# Antimicrobial resistance and comparative genome analysis of high-risk *Escherichia coli* clones isolated from Egyptian children with diarrhoea

**DOI:** 10.1101/2025.10.02.680073

**Authors:** Radwa Abdelwahab, Munirah M. Alhammadi, Muhammad Yasir, Ehsan A. Hassan, Entsar H. Ahmed, Nagla H. Abu-Faddan, Enas A. Daef, Stephen J. W. Busby, Douglas F. Browning

**Affiliations:** Institute of Microbiology and Infection, School of Biosciences, University of Birmingham, Birmingham, B15 2TT, UK; Faculty of Medicine, Assiut University, Assiut 71515, Egypt; Department of Biology, College of Science, Princess Nourah bint Abdulrahman University, P.O. Box 84428, Riyadh 11671, Saudi Arabia; Quadram Institute Bioscience, Norwich Research Park, Norwich, NR4 7UQ, UK; College of Health and Life Sciences, Aston University, Aston Triangle, Birmingham, B4 7ET, UK

**Keywords:** *Escherichia coli*, antibiotic resistance, carbapenemase, virulence, plasmids, whole genome sequencing

## Abstract

*Escherichia coli* is an important human pathogen that is able to cause a variety of infections, which can result in diarrhoea, urinary tract infections, sepsis, and even meningitis, depending on the pathotype of the infecting strain. Like many Gram-negative bacteria, *E. coli* is becoming increasingly resistant to many frontline antibiotics, including third generation cephalosporins and carbapenems, which are often considered the antibiotics of last resort for these infections. This is particularly the case in Egypt, where multidrug resistance (MDR) *E. coli* is highly prevalent. However, in spite of this, few Egyptian MDR *E. coli* strains have been fully characterised by genome sequencing. Here, we present the genome sequences of ten highly MDR *E. coli* strains, which were isolated from children, who presented with diarrhoea at the Outpatients Clinic of Assiut University Children's Hospital in Assiut, Egypt. We report that they carry multiple antimicrobial resistance genes, which includes extended spectrum β-lactamase genes, as well as *bla*NDM and *bla*OXA carbapenemase genes, encoded on IncX3 and potentially IncF plasmids. Many of these stains were also found to be high-risk extra-intestinal pathogenic *E. coli* (ExPEC) clones, belonging to sequence types ST167, ST410 and ST617. Thus, their presence in the Egyptian paediatric population is particularly worrying, and this highlights the need for increased surveillance of high-priority pathogens in this part of the world.

## Introduction

The Gram-negative bacterium *Escherichia coli* is often considered part of the normal gut flora of warm-blooded vertebrate animals, where it acts as commensal organism [1, 2]. However, due to the acquisition of specific virulence genes, some *E. coli* strains can cause disease at various sites within the human body [1, 2]. Diarrhoeagenic *E. coli* are important human pathogens, which result in considerable global morbidity and mortality, particularly amongst children in developing countries. These pathogenic strains are grouped into different pathotypes, based on their disease characteristics, the toxins they secrete and their specific adherence patterns, and includes pathotypes such as enteroaggregative *E. coli* (EAEC), enteropathogenic *E. coli* (EPEC) and enterotoxigenic *E. coli* (ETEC), amongst others [1, 2]. Additionally, other *E. coli* strains have acquired specific virulence determinants that enable them to cause extra-intestinal infections, such as urinary tract infections, sepsis, and meningitis, being termed extra-intestinal pathogenic *E. coli* (ExPEC) [1, 2]. Thus, pathogenic *E. coli* strains are able to cause a considerable spectrum of disease, depending on their particular genic makeup.

Like many bacteria, both clinically and environmentally isolated *E. coli* strains are becoming increasingly resistant to many classes of antibiotics, resulting in emergence of multidrug resistant (MDR) strains. This has led the World Health Organisation (WHO) to categories antibiotic-resistant Gram-negative bacteria, such as *E. coli*, as high-priority pathogens [3]. In particular, this includes *E. coli* strains that are resistant to third-generation cephalosporin antibiotics (*e.g*., ceftriaxone), due to the presence of extended-spectrum β-lactamases (ESBLs), and resistance to carbapenem antibiotics (e.g., imipenem and meropenem), which are often the antibiotics of last resort for many bacterial infections [3-5]. Most Carbapenem-Resistant *Enterobacteriaceae* (CRE) carry Ambler class A, B or D carbapenemases, which includes the *Klebsiella pneumoniae* carbapenemase (KPC), New Delhi metallo-β-lactamase (NDM) and oxacillin hydrolyzing enzymes (OXA-48-like), respectively [4, 5]. These resistance genes are often encoded on mobile genetic elements, such as transposons, integrons and conjugative plasmids, which facilitate their spread, frequently in conjunction with other antimicrobial resistance genes (AMGs). Thus, the global prevalence of CRE is causing a significant public health catastrophe as treatment options dwindle [3-6].

This is particularly the case in Egypt, where CRE are extremely prevalent, being detected at high rates [7, 8]. However, despite this, comparatively few Egyptian carbapenem resistant *E. coli* strains have been fully characterised by genome sequencing. Of those that have, it is clear that they often carry multiple ESBLs (*e.g*., *bla*CTX-M-15 and *bla*TEM-1B) and carbapenemase genes (*e.g*., *bla*NDM-1, *bla*NDM-5, *bla*OXA-48 and *bla*OXA-181), making them resistant to β-lactam antibiotics [8-15]. In many instances, carbapenemase genes are located on conjugative plasmids, co-localised with other AMGs, facilitating their transfer and spread [10, 11]. In a previous study, we isolated a number of *E. coli* strains from children (aged from 2 months to 5 years old), who presented with diarrhoea at the Outpatients Clinic of Assiut University Children's Hospital [15]. Troublingly, many of these isolates are resistant to all tested antibiotics, including carbapenems. As patients had not been previously admitted to hospital, this provided an opportunity to examine the *E. coli* strains circulating within the paediatric population in Assiut. Thus, to understand more about these strains and to identify the AMGs, plasmids and virulence determinants they carry we have characterised 10 of these highly MDR isolates, using whole genome sequencing. We show that the many of the stains are in fact high-risk ExPEC clones, belonging to sequence types ST167, ST410 and ST617 [16-19], and that they possess multiple AMGs, including *bla*NDM and *bla*OXA carbapenemase genes, which are encode on IncX3 and potentially IncF plasmids.

## 2. Materials and Methods

### 2.1. Isolation and characterisation of E. coli strains

The current work is a retrospective study analysing *E. coli* strains collected in 2016 from infants and children (aged from 2 months to 5 years old), who presented with diarrhoea at the Outpatients Clinic of Assiut University Children's Hospital [15]. Ethical approval was granted by the Medical School Ethical Review Board before sample collection proceeded [15]. Individuals from this study had frequent watery diarrhoea (>3 times/ day), with or without blood or mucus, and participants who had received antibiotics within the last 72 hours were excluded from the study. One *E. coli* strain was isolated per patient and identification was carried out at the Medical Research Center, Faculty of Medicine, Assiut University. Resistance to various antimicrobial agents was investigated using the Kirby-Bauer disc diffusion method [20], with interpretation as specified by CLSI 2014 [21]. Antimicrobial discs (Hi-Media, India) contained the following antibiotics: imipenem (10 µg), meropenem (10 µg), cefaclor (30 µg), ceftriaxone (30 µg), amoxicillin (25 µg), ampicillin (10 µg), ciprofloxacin (5 µg), norfloxacin: (10 µg), tobramycin: (10 µg), amikacin (30 µg) oxytetracycline (30 µg) and trimethoprim/ sulfamethoxazole (5 µg) (Supplementary Table S1).

### 2.2. Genome sequencing

The draft genome sequencing of each *E. coli* strain was carried out using Illumina sequencing by Microbes NG (https://microbesng.com/) as detailed previously [15]. Illumina reads were adapter trimmed using Trimmomatic 0.30 with a sliding window quality cutoff of Q15 [22]. Genome assembly was performed using Unicycler v0.4.0 [23] and contigs were annotated using Prokka 1.11 [24]. This Whole Genome Shotgun project has been deposited at DDBJ/ENA/GenBank with the sequence data (BioProject: PRJNA1298299) under the accession numbers: E4: JBQGXB000000000, E15: JBQGXA000000000, E23: JBQGWZ000000000, E27: JBQGWY000000000, E28: JBQGWX000000000, E29: JBQGWW000000000, E30: JBQGWV000000000, E34: JBQGWU000000000, E35: JBQGWT000000000 and E43: JBQGWS000000000.

### 2.3. Bioinformatic analysis of genome sequences

Draft genomes were visualized with Artemis [25], genomes were compared using the Proksee Server (https://proksee.ca/about) [26], the Artemis Comparison Tool (ACT) [27] and the Basic Local Alignment Search Tool (BLAST) at NCBI (https://blast.ncbi.nlm.nih.gov/Blast.cgi). Figures showing genome organization were drawn using ACT [27] and the Proksee Server [26].

Sequence types were determined using MLST 2.0 [28], bacterial serotyping was determined using SerotypeFinder 2.0 [29], plasmids were identified by detecting plasmid replicons using PlasmidFinder 2.1 [30] and virulence gene analysis was performed using VirulenceFinder 2.0 [31, 32] and PathogenFinder 1.1 and 2 [33, 34] using software at the Center for Genomic Epidemiology (CGE) (http://www.genomicepidemiology.org/). Antibiotic resistance genes were detected using ResFinder 3.2 also at CGE [35] (using ResFinder (22/3/2024) and PointFinder (8/3/2024) databases with settings of 90% and 60% for threshold and length ID, respectively). The phylotype of each strain was determined using the EzClermont *in silico* Clermont phylotyper (https://ezclermont.hutton.ac.uk/) [36]. Insertion sequences and bacteriophage were identified using ISfinder (https://www-is.biotoul.fr/index.php) [37] and PHASTER (https://phaster.ca/) [38], respectively.

The phylogenetic analysis of strains was carried out by recreating the phylogenetic tree from Abdelwahab *et al*. [15], using the genomes listed in Figure. 1, and the draft genomes generated in this study. The tree was generated using the AutoMLST2 web server for microbial phylogeny (https://automlst2.ziemertlab.com/) using the standard Denovo Mode with default settings [39]. The phylogenetic tree was visualized, and the *E. coli* branches selected, using the NCBI Tree Viewer (https://www.ncbi.nlm.nih.gov/tools/treeviewer/).

**Figure 1.**
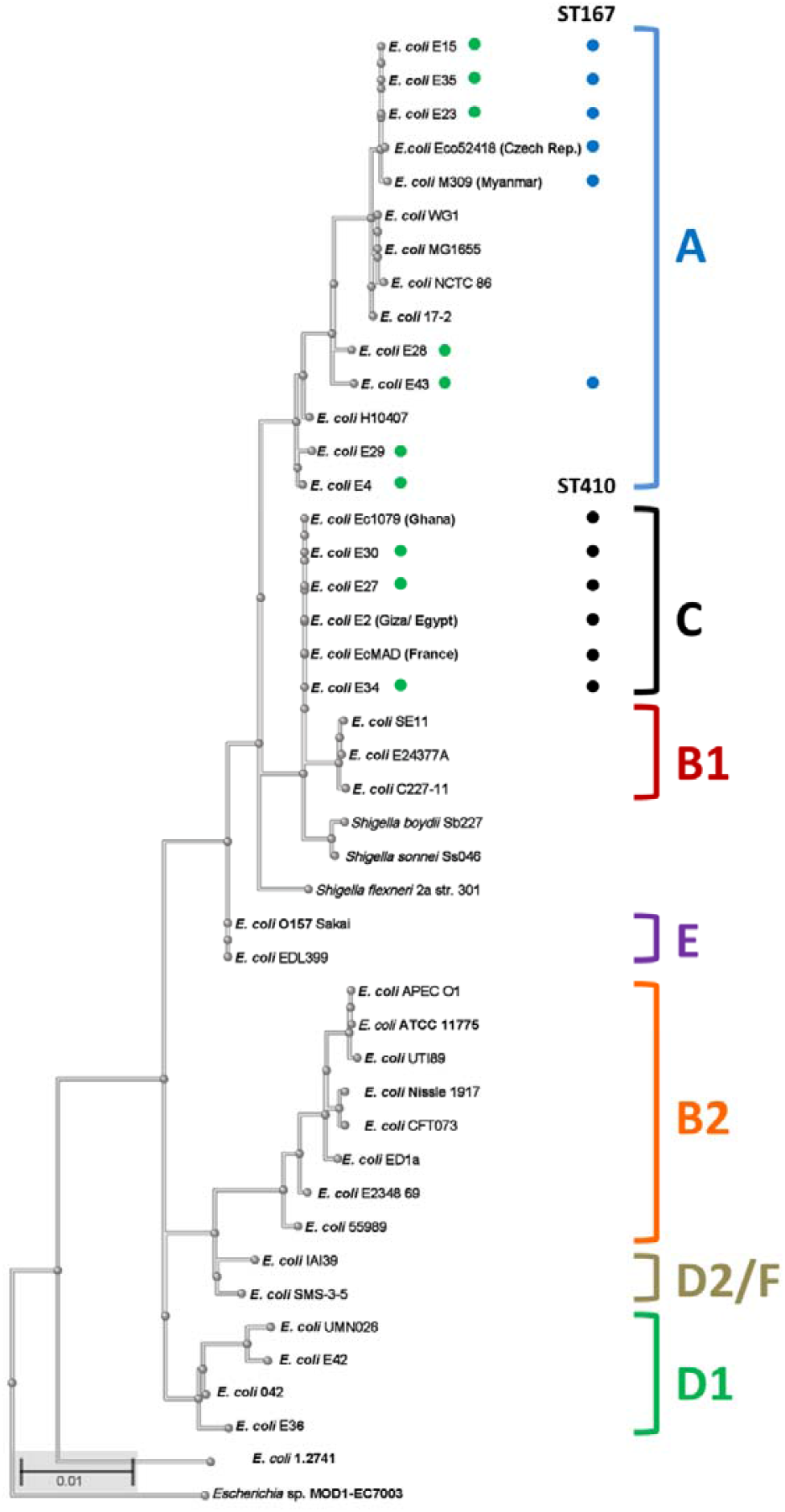
Phylogenetic analysis of the *E. coli* strains isolated in this study. The figure shows a phylogenetic tree of various *E. coli* strains, highlighting the position of the strains investigated in this study (green dots). The tree was reconstructed from the strains in Abdelwahab *et al*. [15], using AutoMLST2 (https://automlst2.ziemertlab.com/) [39]. The various *E. coli* phylotypes are indicated and sequence types ST167 and ST410 are indicated by blue and black dots, respectively.

## 3. Results

### 3.1. Isolation and genome characterisation of E. coli strains

Previously, we isolated 50 *E. coli* strains from children with diarrhoea at the Outpatients Clinic of Assiut University Children's Hospital [15]. To determine their AMR profile strains were tested against a range of frontline antimicrobial agents including carbapenems (imipenem and meropenem), cephalosporins (ceftriaxone and cefaclor), penicillins (amoxicillin and ampicillin), fluoroquinolones (ciprofloxacin and norfloxacin), aminoglycosides (tobramycin and amikacin), tetracycline and folate pathway inhibitors (trimethoprim/ sulphamethoxazole). Many of the *E. coli* strains possessed resistance to all agents tested (Table 1; Supplementary Table S1), highlighting the MDR phenotype of these strains [40].

**Table 1.**
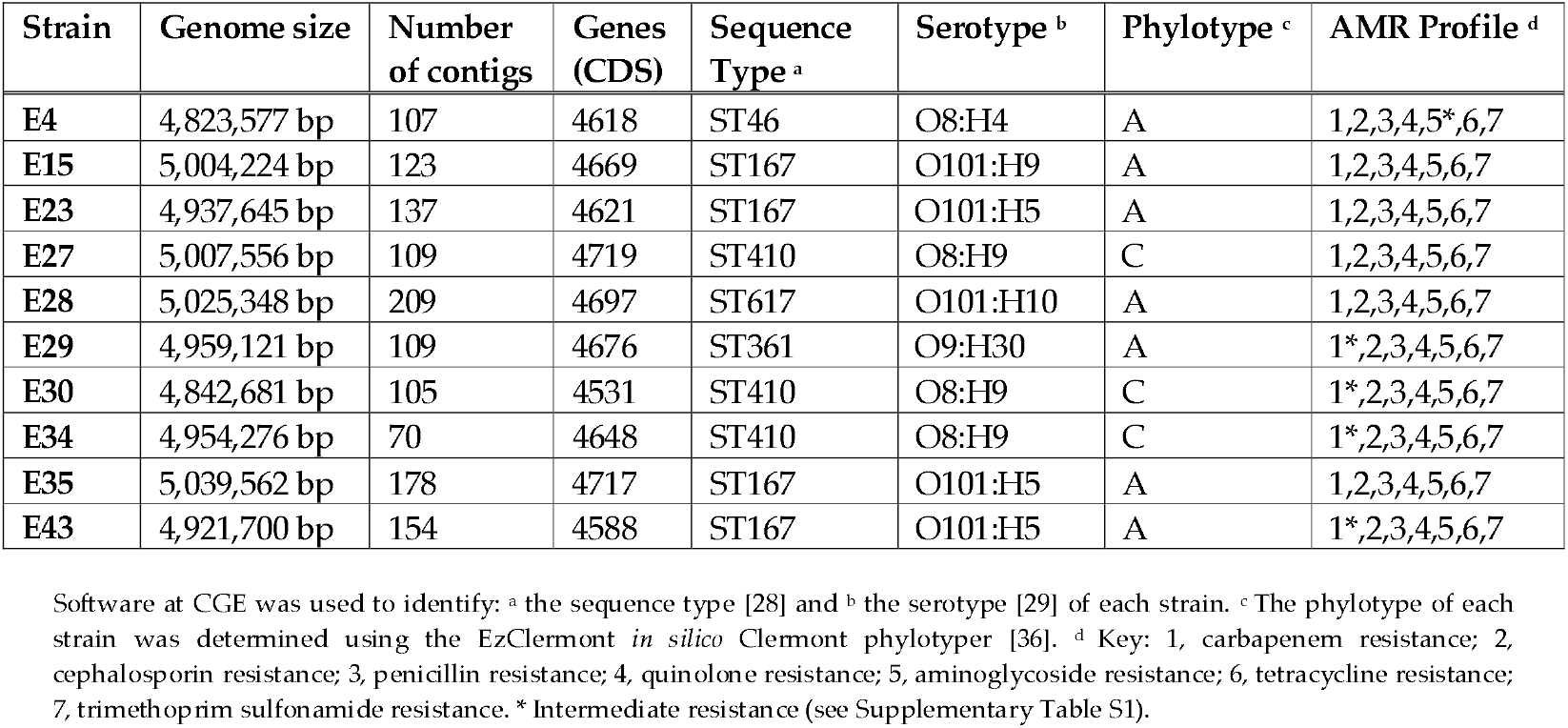
Analysis of the draft genomes and AMR profiles of the *E. coli* strains isolated in this study.

To understand more about the plasmids, AMGs and virulence determinants that each strain possessed, the genomes of the ten most resistant strains were sequenced using short-read Illumina whole genome sequencing (Table 1). Phylogenetic analysis of these MDR strains indicated that four isolates were sequence type ST167, three were ST410 and three were ST46, ST617 and ST361 (Figure 1; Table 1). It is of note that ST167 and ST410 have become global high-risk clones, harbouring multiple AMGs, thus, their presence in the Egyptian population is perhaps not surprising [16-19]. Typical of MDR *E. coli* strains, most strains carried multiple plasmid replicons and, therefore, likely possess a number of different plasmids (Table 2).

**Table 2.**
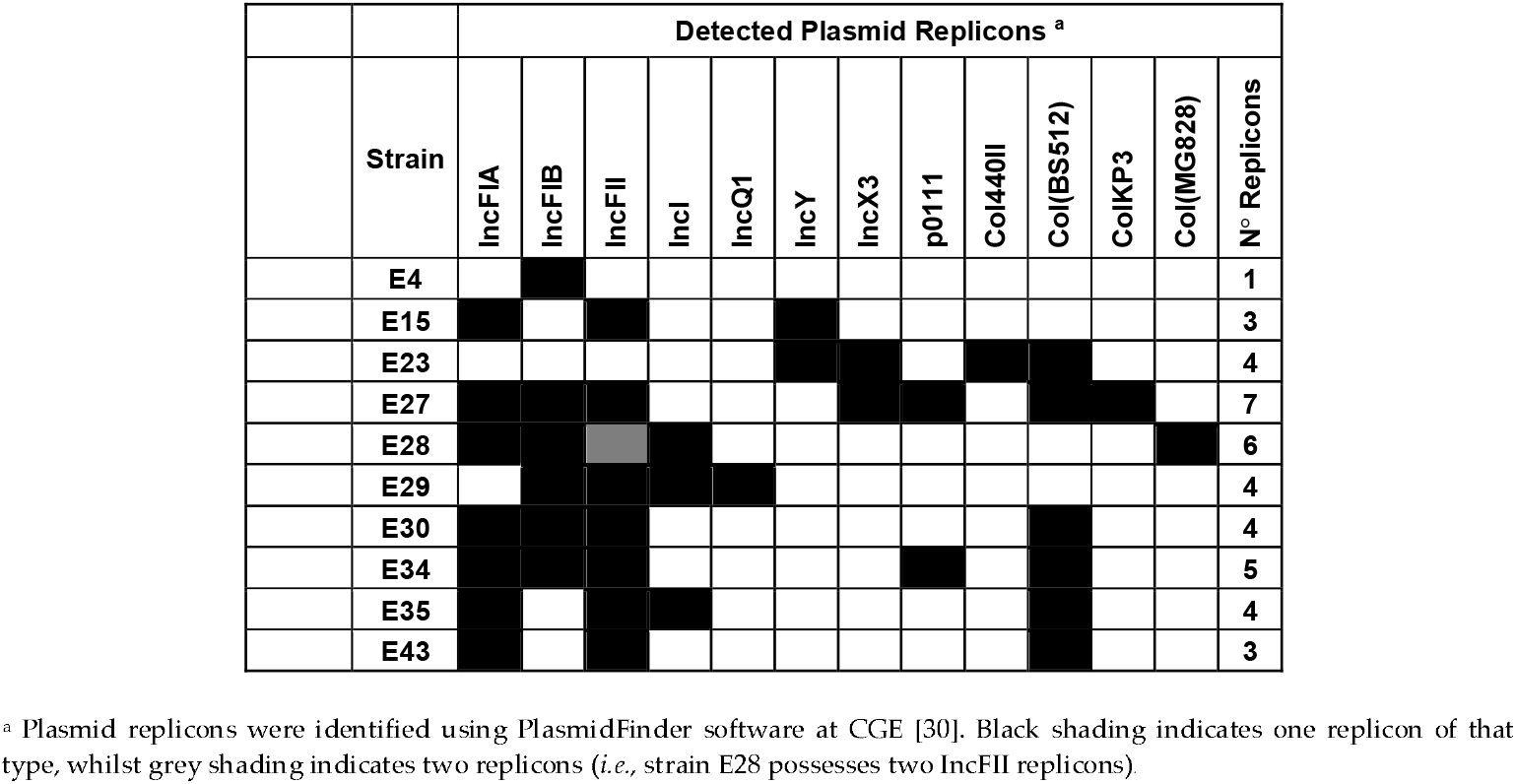
Analysis of plasmid replicons detected in the draft genomes of the *E. coli* strains isolated in this study.

### 3.2. Analysis of acquired AMR genes and chromosomal point mutations associated with AMR

Consistent with their antimicrobial susceptibility pattern (Table 1; Supplementary Table S1) each strain carried multiple AMGs (Table 3). All strains possessed genes that would confer resistance to β-lactam antibiotics (*e.g*., *bla*CTX-M-15, *bla*CMY-2, *bla*CMY-42, *bla*OXA-1, *bla*OXA-9 and *bla*TEM-1B) with the ESBL genes *bla*CTX-M-15 and *bla*TEM-1B each found in 8/10 draft genomes (Table 3). Importantly, genes encoding carbapenemases (*bla*NDM-1, *bla*NDM-5, *bla*NDM-19, *bla*OXA-181 and *bla*OXA-244) were found in six isolates (*i.e*., E4, E15, E23, E27, E35 and E43) explaining these strains resistance to the carbapenem antibiotics, imipenem and meropenem (Table 3; Supplementary Table S1). Furthermore, genes were detected that would result in resistance to fluoroquinolone (*qnrS1* and *aac*(6')-Ib-cr), aminoglycoside (*aac*(3)-IId, *aac*(6')-Ib-cr, *aadA1, aadA2, aadA5, aph*(3')-Ia, *aph*(3'')-Ib, *aph*(6)-Id and *rmtB*), macrolide (*mphA*), sulphonamide (*sul1* and *sul2*), trimethoprim (*dfrA1, dfrA12, dfrA14* and *dfrA17*), tetracycline (*tetA* and tetB) and chloramphenicol (catB3) antibiotics (Table 3). In addition, most strains possessed chromosomal point mutation in *gyrA, parC* and *parE* that are associated with quinolone resistance (*i.e*., resistance to nalidixic acid and ciprofloxacin) [35] (Supplementary Table S2). Coupled with this, 9/10 isolates also possessed either *qnrS1* and/or *aac*(6’)-Ib-cr genes, explaining the complete resistance of all strains to ciprofloxacin. Similarly, 9/10 strains possessed *tet* resistance genes explaining the high level of tetracycline resistance observed (Tables 1 and 3; Supplementary Table S1). Thus, the identification of these resistance determinants correlates with the MDR phenotype observed for these 10 isolates.

**Table 3.**
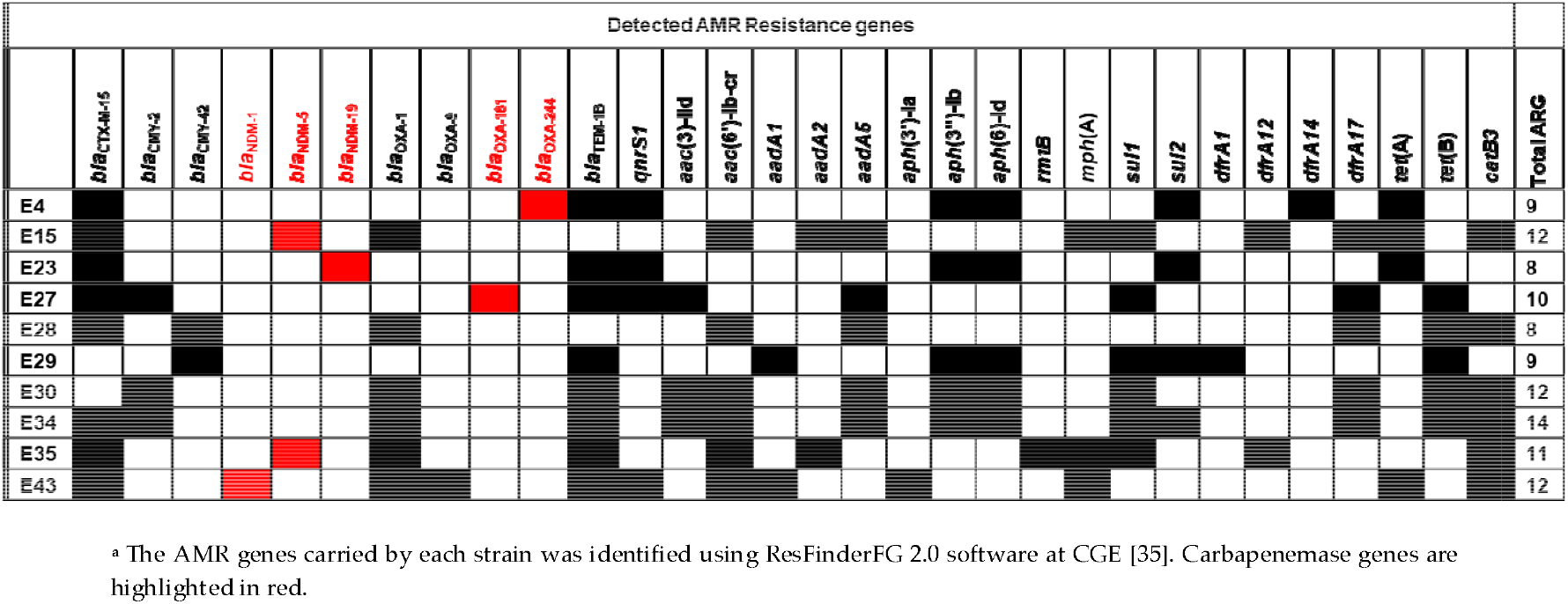
Analysis of AMR genes detected in the draft genomes of *E. coli* strains isolated from children at Assiut Children’s Hospital Outpatients Clinic.

### 3.3. Characterisation of IncX3 plasmids carrying carbapenem resistance determinants

As carbapenem antibiotics are considered the last line of defence against many bacterial pathogens, we sought to understand more about the carbapenemase genes that our strains carry, in particular focusing on the plasmids that might harbour them and lead to their dissemination. Analysis indicated that isolate E23 carries both the *bla*NDM-19 carbapenemase gene and the IncX3 plasmid replicon on a single large contig (contig 27: 46,073 bp), suggesting that this might represent a complete plasmid (Tables 2 and 3). BLAST analysis indicated that this contig was identical (100% coverage: 100% identity) to plasmid pLAU-NDM19 (CP074195.1: 47,332 bp), which was isolated in Lebanon from *E. coli* strain EC20 in 2018 [41] (Figure 2A: Supplementary Figure S1). Plasmid pLAU-NDM19 was shown to be conjugative and strain EC20 was also sequence type ST167. Thus, we propose that strain E23 carries a similar *bla*NDM-19-encoding IncX3 plasmid, which we have termed pE23-NDM19 (Figure 2A).

**Figure 2.**
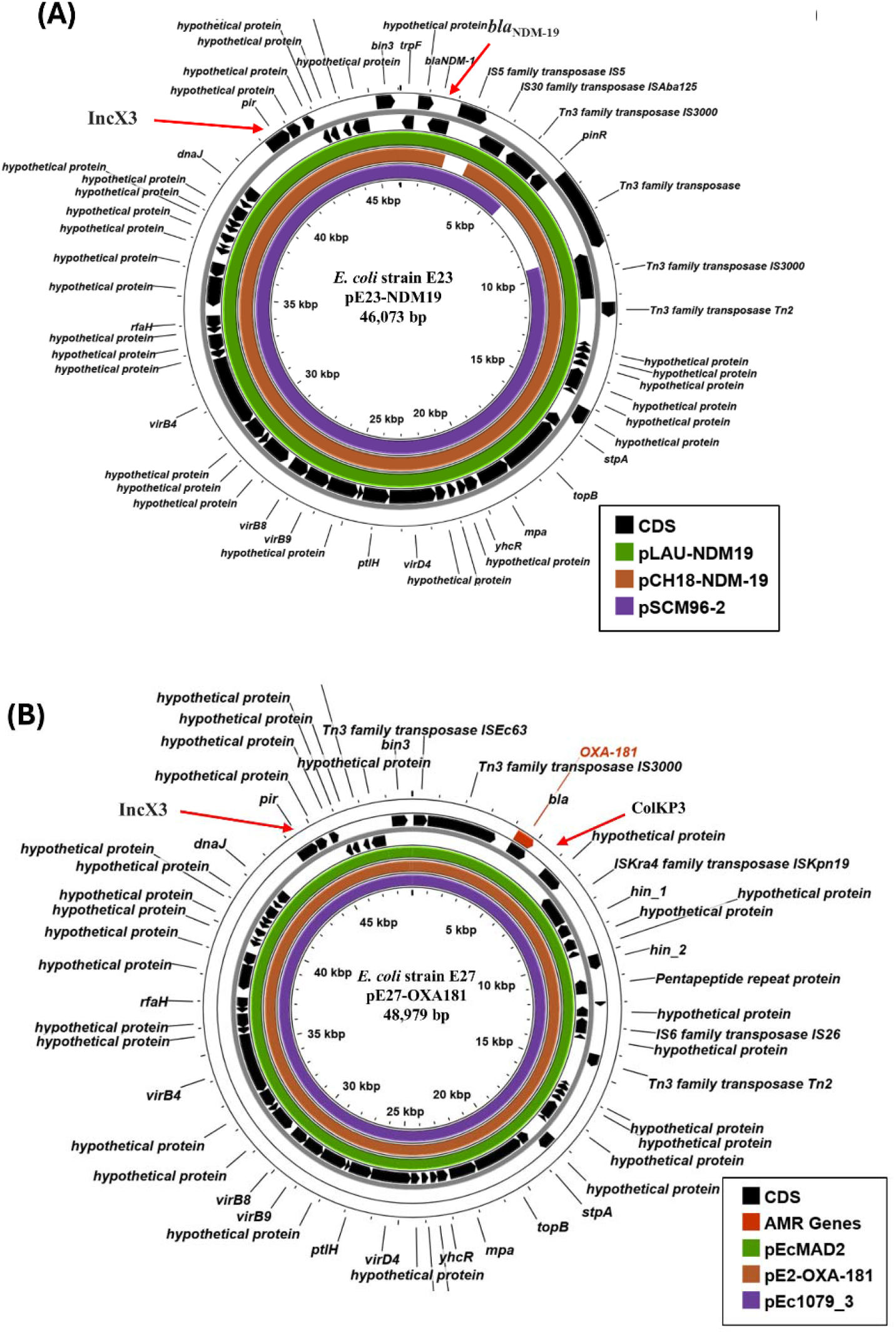
Analysis of the IncX3 plasmids carried by *E. coli* isolates E23 and E27. (A) The panel shows the comparison of pE23-NDM19 (E23 contig 27: 46,073 bp) with plasmids pLAU-NDM19 (CP074195.1; 47,332 bp) [41], pCH18-NDM-19 (MK091521: 48,737 bp) and pSCM96-2 (CP028718.1: 46,161 bp) using ProkSee [26]. The genes (CDS) of pE23-NDM19 are displayed in the outer rings, with the location of *bla*NDM-19 and the IncX3 replicon indicated. The green, brown and purple rings depict the BLAST results when the sequences of pLAU-NDM19, pCH18-NDM-19 and pSCM96-2 are compared with pE23-NDM19. (B) The panel shows the comparison of pE27-OXA181 (E27 contig 22: 48,979 bp) with plasmids pEcMAD2 (LR595693.1: 51,479 bp) [42], pE2-OXA-181 (CP048918.1: 51,479 bp) [9] and pEc1079_3 (CP081309.1: 51,479 bp) [43] using ProkSee [26]. The genes (CDS) of pE27-OXA181 are displayed in the outer rings, with the location of *bla*OXA-181 and the IncX3 and ColKP3 replicons indicated. The green, brown and purple rings depict the BLAST results when the sequences of pEcMAD2, pE2-OXA-181 and pEc1079_3 are compared with pE27-OXA181.

Out of all our strains, E27 carries the most plasmid replicons, possessing seven in total (Table 2). In this instance, the *bla*OXA-181 carbapenemase gene co-localises with the IncX3 and ColKP3 replicons on a single large contig (contig 22: 48,979 bp) (Figure 2B). Analysis indicated this contig was identical (100% coverage: 100% identity) to plasmid pE2-OXA-181 (CP048918.1: 51,479 bp) isolated in Egypt (Giza) in 2015 [9], plasmid pEcMAD2 (LR595693.1: 51,479 bp) isolated in France in 2013 [42], and pEc1079_3 (CP081309.1: 51,479 bp) form Ghana in 2015 [43] (Figure 2B; Supplementary Figure S2). Thus, strain E27 carries a similar *bla*OXA-181-encoding IncX3-ColKP3 plasmid, which we term pE27-OXA181 (Figure 2B). Interestingly, *E. coli* strains E2, EcMAD1, and Ec1079 from which these plasmids came were sequence type ST410, like E27, and possess very similar chromosomes with only a few minor regions of difference (Figure 1; Supplementary Figure S3; Table 1) [9, 42, 43].

### 3.4. Characterisation of IncF plasmids potentially carrying carbapenem resistance genes

Due to the limitations of short-read sequencing, the carbapenemase genes carried by other strains were not co-localised on contigs that possessed plasmid replicons. However, in some instances, we are able to make suggestions concerning the plasmids that might harbour these genes. Our analysis indicated that for isolate E15 (sequence type ST167), the *bla*NDM-5 gene was located on a small contig (contig 53: 3,163 bp), whilst E15 carried three plasmid replicons (Table 2). BLASTn analysis of the IncFII (contig 38: 16,561 bp) and the IncFIA (contig 40: 14,118bp) replicons indicated that they were very similar to sections of plasmid p52148_NDM5 (CP050384.1: 121,872 bp), which also carries *bla*NDM-5 (Figure 3A) (100%/ 99% coverage: 99.99%/ 99.96% identity, respectively). Strain Eco52148 was isolated in 2019 from a patient who was repatriated from northern Africa to the Czech Republic and, like E15, was sequence type ST167 (Figure 1) [44]. As both p52148_NDM5 and E15 carry *mphA, sul1, aadA2, dfrA12* and *tetA* AMGs (Table 3; Figure 3A), we propose that E15 carries a similar IncFII-IncFIA resistance plasmid.

**Figure 3.**
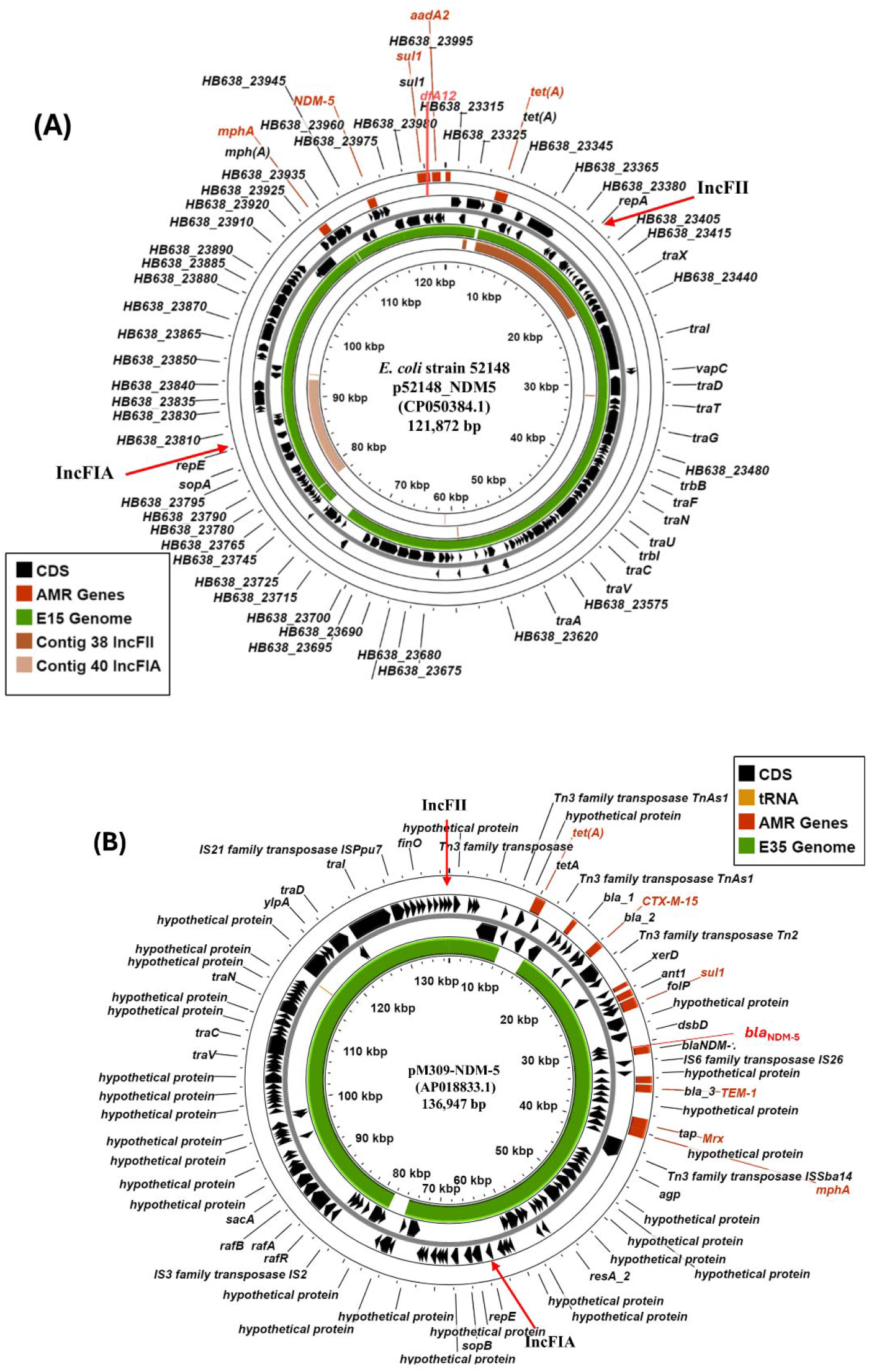
Analysis of IncF plasmids carrying *bla*NDM-5 carbapenemase genes in *E. coli* isolates E15 and E35. (A) Comparison of *E. coli* Eco52148 plasmid p52148_NDM5 with the draft genome of E15. The panel shows the comparison of p52148_NDM5 (CP050384.1: 121,872 bp) [44] with the draft genome of E15 and E15 contigs 38 (16,561 bp) and 40 (14,118bp), using ProkSee [26]. The outer two rings display the genes of p52148_NDM5 (CDS) on both strands. The green, brown and light brown rings illustrate the BLAST results when the E15 draft genome and contigs 38 and 40, respectively, are compared to p52148_NDM5. The location of the *bla*NDM-5, *mphA, sul1, aadA2, dfrA12* and *tetA* AMR genes and the IncFII and IncFIA plasmid replicons are shown. (B) Comparison of plasmid pM309-NDM5 with the draft genome sequence of *E. coli* isolate E35. The figure shows the comparison of pM309-NDM5 (AP018833.1: 136,947 bp) [45] with the draft genome of E35, using ProkSee [26]. The genes (CDS) of pM309-NDM5 are displayed in the outer rings, with the location of the various AMR genes (including *bla*NDM-5) and plasmid replicons (IncFIA and IncFII) indicated. The green ring depicts the BLAST results when the E35 draft genome is compared with pM309-NDM5.

Strain E35 (also ST167) carries the *bla*NDM-5 carbapenemase gene on contig 38 (38,086 bp), which co-localises with *bla*CTX-M-15, *bla*TEM-1B, *mphA, rmtB, aadA2, sul1* and *dfrA12* AMGs. BLASTn analysis indicated that this contig was identical (100% coverage: 100% identity) to plasmid pM309-NDM5, carried by ST167 *E. coli* strain M309 (AP018833.1: 136,947 bp), isolated from Yangon, Myanmar in 2015 (Figures 1 and 3B; Supplementary Figure S4) [45]. Furthermore, E35 contigs carrying the IncFIA (contig 45: 15,794 bp) and IncFII (contig 49: 11,728 bp) replicons were very similar to those carried by pM309-NDM5 (100%/ 98% coverage: 100%/ 100%, respectively) (Supplementary Figure S4). Thus, we propose that isolate E35 carries a large dual IncFIA-IncFII plasmid, which likely encodes *bla*NDM-5.

### 3.4. Carriage of virulence-associated genes in Egyptian E. coli isolates

The analysis of each strain using PathogenFinder indicated that all were potential human pathogens, carrying a number of characterized virulence determinants (Supplementary Table S3) [31-34]. This included glutamate decarboxylases genes (*gadA*/ *gadB*) involved in acid resistance [46], *iss/ bor*, which encodes a lipoprotein involved in increased serum survival [47], *capU*, a hexosyltransferase homologue [48], *hra* heat-resistant agglutinin [49], *traT*, an outer membrane protein is involved in serum resistance [50], and *lpfA* long polar fimbriae [49]. In addition, both E15 and E28 possesses the yersiniabactin siderophore uptake system (*ipr2/ fyuA*) with E28 carrying additional iron scavenging systems (*i.e*., *sitA* iron transport protein and the *iucC*/*iutA* aerobactin system) [51]. It is of note, the carriage of specific virulence genes was sequence type-specific, with ST167 strains (E15, E23, E35 and E43) and ST410 strains (E27, E30 and E34), possessing similar respective virulence profiles (Supplementary Table S3). For ST167 strains many of the virulence genes they carry (*e.g*., *iss, hra, traT and ipr2/ fyuA*) are associated with ExPEC [52-54] to which sequence type ST167 has been grouped [16]. Interestingly, for E28 (ExPEC sequence type ST617 [16-19]), *sitABCD* and the aerobactin siderophore gene cluster (*iucABC-iutA*) are located on contig 36 (43,206 bp) flanked by IncFIA and IncFIB replicons, suggesting they are plasmid borne (Supplementary Figure S5). Analysis of this contig and E28 contig 22 (75,303 bp) indicated that they were similar (100%/ 100% coverage: 100/ 99.96% identity, respectively) to sections of pEC22-OXA-1 (CP084902.1: 169208 bp) isolated from *E. coli* strain Ec20 in Jinhua, China in 2019. Like pEC22-OXA-1, contig 22 carries *aac(*6’)-Ib-cr, *aadA5, bla*CTX-M-15, *bla*OXA-1, *dfrA17*, two IncFII plasmid replicons and numerous *tra* genes (Supplementary Figure S5), suggesting that E28 might also carry a hybrid virulence/ AMR plasmid, potentially capable of conjugative transfer.

## 4. Discussion

In this study, we have characterized ten MDR *E. coli* strains isolated from infants and children with diarrhoea that attended the Outpatients Clinic of Assiut University Children’s Hospital in 2016 [15]. It is of note that none of our strains carried virulence determinants that are associated with diarrhoeagenic *Escherichia coli* pathotypes, such as EAEC, EPEC or ETEC [1]. Rather, they possessed virulence genes associated with the ExPEC pathotype (Supplementary Table S3) with many strains belonging to ExPEC-associated sequence types (*i.e*., ST167, ST410 and ST617) [16-19]. Thus, we think it unlikely that the strains we have isolated are the cause of diarrhoea in these individuals. As the human alimentary canal is a considered a reservoir for some ExPEC infections [16, 55], it is particularly alarming that carbapenem resistant ExPEC strains were circulating in the paediatric population in Egypt at this time.

All strains were resistant to multiple antibiotics and carried numerous AMGs (Tables 1 and 3; Supplementary Table 1), with *bla*NDM or *bla*OXA carbapenemase genes detected in six isolates. Thus, treatment options for infections associated with these strains would likely be limited, however, genome analysis suggests that they all remain sensitive to fosfomycin, tigecycline and colistin antibiotics. Four of these CRE strains (*i.e*., E15, E23, E35 and E43) are considered high-risk ExPEC clones, being sequence type ST167, and all carried variants of the *bla*NDM carbapenemase (Table 3) [16, 17, 19]. It is of note that the *bla*NDM-1 and *bla*NDM-5 alleles have been detected in Egyptian ST167 strains before [8, 56]. In spite of the limitations of short-read genome sequencing, we propose that the *bla*NDM genes from strains E15 and E35 may be carried on large IncFIA-IncFII dual replicon plasmids, which is a common occurrence (Figure 3) [4, 44, 45, 56]. Conversely, for strain E23 the *bla*NDM-19 carbapenemase gene was encoded on an IncX3 plasmid, pE23-NDM19 (Figure 2A). The NDM-19 carbapenemase was first characterised in 2019, differing from NDM-1 by three amino acid substitutions (*i.e*., D130N, M154L and A233V) (Supplementary Figure 1) and confers high-level resistance to 3rd generation cephalosporins and carbapenems under zinc-limited conditions, which are thought to prevail at infections sites [57, 58]. Thus, its appearance is an important escalation in NDM evolution [58]. Like pE23-NDM19, early isolates carried the *bla*NDM-19 gene on conjugational IncX3 plasmids, *e.g*. pSCM96-2 (CP028718.1: 46,161 bp) which was isolated in China in 2017 and pCH18-NDM-19 (MK091521: 48,737 bp), which was isolated from an Egyptian patient in Switzerland in 2018 [57, 58] (Figure 2A). As our strains predate the isolation of these first plasmids, it is clear that IncX3/ *bla*NDM-19 plasmids were already present within the Egyptian population as early as 2016.

Three of our strains were sequence type ST410 (*i.e*., E27, E30 and E34), which is also considered a high-risk ExPEC clone [16, 18, 19]. Strain E27, carries the *bla*OXA-181 carbapenemase on a dual IncX3-ColKP3 plasmid pE27-OXA181, which was similar to plasmids isolated from ST410 strains in France, Egypt and Ghana (*i.e*., pEcMAD2, pE2-OXA-181 and pEc1079_3) (Figures 1 and 2B; Supplementary Figure S2) [9, 42, 43]. Additionally, the chromosomes of these strains (*i.e*., E2, EcMAD1, and Ec1079) are very similar to those of E27, E30 and E34, with only a few minor regions of difference (Supplementary Figure S3). Our analysis also suggests that these strains likely share additional plasmids. For example, both the E27 and E34 carry a p0111 replicon (contigs 18: 92,130 bp and 16: 96,506 bp, respectively) (Table 2), which is also on plasmid pE2-2 carried by strain E2 (Giza/ Egypt) (CP048917.1: 92,027 bp) [9] (Supplementary Figure S6). Furthermore, strains EcMAD1, E2, and Ec1079 also carry an IncFIA/IncFIB/IncFII multi-replicon plasmid (*i.e*., pEcMAD1, pE2-NDM-CTX-M and pEc1079_1, respectively) that could be detected in the draft genomes of our three ST410 strains (Supplementary Figure S7). Thus, there seems to be a close relationship between these ST140 strains and the plasmids that they carry (Figure 1).

In this work, we have been able to gain a snapshot of the MDR *E. coli* strains present in the paediatric population in Assiut in 2016. It is particularly concerning that, due to the lack of funding and routine sequencing in Egypt [59-61], it has taken considerable time for us to uncover that carbapenem resistant high-risk ExPEC clones were present within the population at this time. As antibiotics are available to purchase in Egypt without prescription [62-64] and there is evidence of unnecessary prescribing, particularly of β-lactams [63, 65], it is likely that this situation could continue to worsen. Coupled with this, the increase in AMR resistance associated with humanitarian disasters in nearby places, such as Gaza, will continue to place a considerable burden on resources [66-68]. Thus, it is clear that there is a need for improved antimicrobial stewardship, infection control and better surveillance of high-priority Gram-negative CREs to combat their spread in, and from, this geographical area.

## Author Contributions

Conceptualization, E.A.H., E.H.A., N.H.A., E.A.D., S.J.W.B and D.F.B.; formal analysis, R.A., M.Y. and D.F.B.; investigation, R.A., M.M.A., M.Y., and D.F.B.; data curation, D.F.B.; writing—original draft preparation, D.F.B.; writing—review and editing, R.A., M.M.A., M.Y., E.A.H., E.H.A., N.H.A., E.A.D., S.J.W.B and D.F.B.; supervision, E.A.D., S.J.W.B. and D.F.B. All authors have read and agreed to the published version of the manuscript.

## Institutional Review Board Statement

The study was conducted in accordance with the Declaration of Helsinki and approved by the Assiut Faculty of Medicine Institutional Review Board of 04-2025-100401 (24/5/2016).

## Informed Consent Statement

As stipulated by the Assiut Faculty of Medicine Institutional Review Board Parental/Gurdian Consent was taken for each sample with elaboration that the stool sample be used to detect and isolate the causative organism, determine its antibiotic sensitivity and investigate its properties for research purposes only. No identifiable data was collected to link patients to any of the samples.

## Data Availability Statement

This Whole Genome Shotgun project has been deposited at DDBJ/ENA/GenBank with the sequence data for E. coli strains (BioProject: PRJNA1298299) under the accession numbers: E4: JBQGXB000000000, E15: JBQGXA000000000, E23: JBQGWZ000000000, E27: JBQGWY000000000, E28: JBQGWX000000000, E29: JBQGWW000000000, E30: JBQGWV000000000, E34: JBQGWU000000000, E35: JBQGWT000000000 and E43: JBQGWS000000000.

## Acknowledgments

We thank MicrobesNG for sequencing and genome annotation.

## References

[1] J.B. Kaper, J.P. Nataro, H.L. Mobley, Pathogenic Escherichia coli, Nat Rev Microbiol. 2 (2004) 123–40. doi: 10.1038/nrmicro818.

[2] L.W. Riley, Distinguishing Pathovars from Nonpathovars: Escherichia coli, Microbiol Spectr. 8 (2020). doi: 10.1128/microbiolspec.AME-0014-2020.

[3] H. Sati, E. Carrara, A. Savoldi, P. Hansen, J. Garlasco, E. Campagnaro, et al., The WHO Bacterial Priority Pathogens List 2024: a prioritisation study to guide research, development, and public health strategies against antimicrobial resistance, Lancet Infect Dis. (2025). doi: 10.1016/s1473-3099(25)00118-5.

[4] W. Wu, Y. Feng, G. Tang, F. Qiao, A. McNally, Z. Zong, NDM Metallo-β-Lactamases and Their Bacterial Producers in Health Care Settings, Clin Microbiol Rev. 32 (2019). doi: 10.1128/cmr.00115-18.

[5] K. Kopotsa, J. Osei Sekyere, N.M. Mbelle, Plasmid evolution in carbapenemase-producing Enterobacteriaceae: a review, Ann N Y Acad Sci. 1457 (2019) 61–91. doi: 10.1111/nyas.14223.

[6] T. Tängdén, C.G. Giske, Global dissemination of extensively drug-resistant carbapenemase-producing Enterobacteriaceae: clinical perspectives on detection, treatment and infection control, J Intern Med. 277 (2015) 501–12. doi: 10.1111/joim.12342.

[7] J.A. Karlowsky, S.H. Lob, K.M. Kazmierczak, R.E. Badal, K. Young, M.R. Motyl, et al., In Vitro Activity of Imipenem against Carbapenemase-Positive Enterobacteriaceae Isolates Collected by the SMART Global Surveillance Program from 2008 to 2014, J Clin Microbiol. 55 (2017) 1638–49. doi: 10.1128/jcm.02316-16.

[8] G. Peirano, L. Chen, D. Nobrega, T.J. Finn, B.N. Kreiswirth, R. DeVinney, et al., Genomic Epidemiology of Global Carbapenemase-Producing Escherichia coli, 2015-2017, Emerg Infect Dis. 28 (2022) 924–31. doi: 10.3201/eid2805.212535.

[9] D. Gamal, M. Fernández-Martínez, I. El-Defrawy, A.A. Ocampo-Sosa, L. Martínez-Martínez, First identification of NDM-5 associated with OXA-181 in Escherichia coli from Egypt, Emerg Microbes Infect. 5 (2016) e30. doi: 10.1038/emi.2016.24.

[10] A.M. Soliman, H. Ramadan, M. Sadek, H. Nariya, T. Shimamoto, L.M. Hiott, et al., Draft genome sequence of a bla(NDM-1)- and bla(OXA-244)-carrying multidrug-resistant Escherichia coli D-ST69 clinical isolate from Egypt, J Glob Antimicrob Resist. 22 (2020) 832–4. doi: 10.1016/j.jgar.2020.07.015.

[11] N.M. Mohamed, A.S. Zakaria, E.A. Edward, Genomic Characterization of International High-Risk Clone ST410 Escherichia coli Co-Harboring ESBL-Encoding Genes and bla(NDM-5) on IncFIA/IncFIB/IncFII/IncQ1 Multireplicon Plasmid and Carrying a Chromosome-Borne bla(CMY-2) from Egypt, Antibiotics (Basel). 11 (2022). doi: 10.3390/antibiotics11081031.

[12] S.D. Braun, S. Rezk, C. Brandt, M. Reinicke, C. Diezel, E. Müller, et al., Tracking Multidrug Resistance in Gram-Negative Bacteria in Alexandria, Egypt (2020-2023): An Integrated Analysis of Patient Data and Diagnostic Tools, Antibiotics (Basel). 13 (2024). doi: 10.3390/antibiotics13121185.

[13] A.S. Zakaria, E.A. Edward, N.M. Mohamed, Pathogenicity Islands in Uropathogenic Escherichia coli Clinical Isolate of the Globally Disseminated O25:H4-ST131 Pandemic Clonal Lineage: First Report from Egypt, Antibiotics (Basel). 11 (2022). doi: 10.3390/antibiotics11111620.

[14] A.M. Soliman, H. Ramadan, T. Shimamoto, T. Komatsu, F. Maruyama, T. Shimamoto, Detection and Genomic Characteristics of NDM-19- and QnrS11-Producing O101:H5 Escherichia coli Strain Phylogroup A: ST167 from a Poultry Farm in Egypt, Microorganisms. 13 (2025). doi: 10.3390/microorganisms13081769.

[15] R. Abdelwahab, M. Yasir, R.E. Godfrey, G.S. Christie, S.J. Element, F. Saville, et al., Antimicrobial resistance and gene regulation in Enteroaggregative Escherichia coli from Egyptian children with diarrhoea: Similarities and differences, Virulence. 12 (2021) 57–74. doi: 10.1080/21505594.2020.1859852.

[16] A.R. Manges, H.M. Geum, A. Guo, T.J. Edens, C.D. Fibke, J.D.D. Pitout, Global Extraintestinal Pathogenic Escherichia coli (ExPEC) Lineages, Clin Microbiol Rev. 32 (2019). doi: 10.1128/cmr.00135-18.

[17] A. Garcia-Fernandez, L. Villa, G. Bibbolino, A. Bressan, M. Trancassini, V. Pietropaolo, et al., Novel Insights and Features of the NDM-5-Producing Escherichia coli Sequence Type 167 High-Risk Clone, mSphere. 5 (2020). doi: 10.1128/mSphere.00269-20.

[18] L. Roer, S. Overballe-Petersen, F. Hansen, K. Schønning, M. Wang, B.L. Røder, et al., Escherichia coli Sequence Type 410 Is Causing New International High-Risk Clones, mSphere. 3 (2018). doi: 10.1128/mSphere.00337-18.

[19] J. Huang, C. Lv, M. Li, T. Rahman, Y.F. Chang, X. Guo, et al., Carbapenem-resistant Escherichia coli exhibit diverse spatiotemporal epidemiological characteristics across the globe, Commun Biol. 7 (2024) 51. doi: 10.1038/s42003-023-05745-7.

[20] A.W. Bauer, W.M. Kirby, J.C. Sherris, M. Turck, Antibiotic susceptibility testing by a standardized single disk method, Am J Clin Pathol. 45 (1966) 493–6.

[21] Clinical_and_Laboratory_Standards_Institute_(CLSI), Performance Standards for Antimicrobial Susceptibility Testing; Twenty-Fourth Informational Supplement, CLSI Document M100-S24, Wayne. 34(1) (2014).

[22] A.M. Bolger, M. Lohse, B. Usadel, Trimmomatic: a flexible trimmer for Illumina sequence data, Bioinformatics. 30 (2014) 2114–20. doi: 10.1093/bioinformatics/btu170.

[23] R.R. Wick, L.M. Judd, C.L. Gorrie, K.E. Holt, Unicycler: Resolving bacterial genome assemblies from short and long sequencing reads, PLoS Comput Biol. 13 (2017) e1005595. doi: 10.1371/journal.pcbi.1005595.

[24] T. Seemann, Prokka: rapid prokaryotic genome annotation, Bioinformatics. 30 (2014) 2068–9. doi: 10.1093/bioinformatics/btu153.

[25] K. Rutherford, J. Parkhill, J. Crook, T. Horsnell, P. Rice, M.A. Rajandream, et al., Artemis: sequence visualization and annotation, Bioinformatics. 16 (2000) 944–5. doi: 10.1093/bioinformatics/16.10.944.

[26] J.R. Grant, E. Enns, E. Marinier, A. Mandal, E.K. Herman, C.Y. Chen, et al., Proksee: in-depth characterization and visualization of bacterial genomes, Nucleic Acids Res. 51 (2023) W484–w92. doi: 10.1093/nar/gkad326.

[27] T.J. Carver, K.M. Rutherford, M. Berriman, M.A. Rajandream, B.G. Barrell, J. Parkhill, ACT: the Artemis Comparison Tool, Bioinformatics. 21 (2005) 3422–3. doi: 10.1093/bioinformatics/bti553.

[28] M.V. Larsen, S. Cosentino, S. Rasmussen, C. Friis, H. Hasman, R.L. Marvig, et al., Multilocus sequence typing of total-genome-sequenced bacteria, J Clin Microbiol. 50 (2012) 1355–61. doi: 10.1128/jcm.06094-11.

[29] K.G. Joensen, A.M. Tetzschner, A. Iguchi, F.M. Aarestrup, F. Scheutz, Rapid and Easy In Silico Serotyping of Escherichia coli Isolates by Use of Whole-Genome Sequencing Data, J Clin Microbiol. 53 (2015) 2410–26. doi: 10.1128/jcm.00008-15.

[30] A. Carattoli, E. Zankari, A. Garcia-Fernandez, M. Voldby Larsen, O. Lund, L. Villa, et al., In silico detection and typing of plasmids using PlasmidFinder and plasmid multilocus sequence typing, Antimicrob Agents Chemother. 58 (2014) 3895–903. doi: 10.1128/aac.02412-14.

[31] K.G. Joensen, F. Scheutz, O. Lund, H. Hasman, R.S. Kaas, E.M. Nielsen, et al., Real-time whole-genome sequencing for routine typing, surveillance, and outbreak detection of verotoxigenic Escherichia coli, J Clin Microbiol. 52 (2014) 1501–10. doi: 10.1128/jcm.03617-13.

[32] A.M. Malberg Tetzschner, J.R. Johnson, B.D. Johnston, O. Lund, F. Scheutz, In Silico Genotyping of Escherichia coli Isolates for Extraintestinal Virulence Genes by Use of Whole-Genome Sequencing Data, J Clin Microbiol. 58 (2020). doi: 10.1128/jcm.01269-20.

[33] S. Cosentino, M. Voldby Larsen, F. Møller Aarestrup, O. Lund, PathogenFinder--distinguishing friend from foe using bacterial whole genome sequence data, PLoS One. 8 (2013) e77302. doi: 10.1371/journal.pone.0077302.

[34] A. Ferrer Florensa, J.J. Almagro Armenteros, R.S. Kaas, P.T.L. Conradsen Clausen, H. Nielsen, B. Rost, et al., Whole-genome prediction of bacterial pathogenic capacity on novel bacteria using protein language models, with PathogenFinder2. (2025) 2025.04.12.648497. doi: 10.1101/2025.04.12.648497 %J bioRxiv.

[35] E. Zankari, H. Hasman, S. Cosentino, M. Vestergaard, S. Rasmussen, O. Lund, et al., Identification of acquired antimicrobial resistance genes, J Antimicrob Chemother. 67 (2012) 2640–4. doi: 10.1093/jac/dks261.

[36] N.R. Waters, F. Abram, F. Brennan, A. Holmes, L. Pritchard, Easy phylotyping of Escherichia coli via the EzClermont web app and command-line tool, Access Microbiol. 2 (2020) acmi000143. doi: 10.1099/acmi.0.000143.

[37] P. Siguier, J. Perochon, L. Lestrade, J. Mahillon, M. Chandler, ISfinder: the reference centre for bacterial insertion sequences, Nucleic Acids Res. 34 (2006) D32–6. doi: 10.1093/nar/gkj014.

[38] D. Arndt, J.R. Grant, A. Marcu, T. Sajed, A. Pon, Y. Liang, et al., PHASTER: a better, faster version of the PHAST phage search tool, Nucleic Acids Res. 44 (2016) W16–21. doi: 10.1093/nar/gkw387.

[39] B. Pourmohsenin, A. Wiese, N. Ziemert, AutoMLST2: a web server for phylogeny and microbial taxonomy, Nucleic Acids Res. 53 (2025) W45–w50. doi: 10.1093/nar/gkaf397.

[40] A.P. Magiorakos, A. Srinivasan, R.B. Carey, Y. Carmeli, M.E. Falagas, C.G. Giske, et al., Multidrug-resistant, extensively drug-resistant and pandrug-resistant bacteria: an international expert proposal for interim standard definitions for acquired resistance, Clin Microbiol Infect. 18 (2012) 268–81. doi: 10.1111/j.1469-0691.2011.03570.x.

[41] J. Moussa, E. Nassour, T. Jisr, M. El Chaar, S. Tokajian, Characterization of bla(NDM-19)-producing IncX3 plasmid isolated from carbapenem-resistant Escherichia coli and Klebsiella pneumoniae, Heliyon. 10 (2024) e29642. doi: 10.1016/j.heliyon.2024.e29642.

[42] R. Patiño-Navarrete, I. Rosinski-Chupin, N. Cabanel, L. Gauthier, J. Takissian, J.Y. Madec, et al., Stepwise evolution and convergent recombination underlie the global dissemination of carbapenemase-producing Escherichia coli, Genome Med. 12 (2020) 10. doi: 10.1186/s13073-019-0699-6.

[43] S. Mahazu, I. Prah, A. Ayibieke, W. Sato, T. Hayashi, T. Suzuki, et al., Possible Dissemination of Escherichia coli Sequence Type 410 Closely Related to B4/H24RxC in Ghana, Front Microbiol. 12 (2021) 770130. doi: 10.3389/fmicb.2021.770130.

[44] K. Chudejova, L. Kraftova, V. Mattioni Marchetti, J. Hrabak, C.C. Papagiannitsis, I. Bitar, Genetic Plurality of OXA/NDM-Encoding Features Characterized From Enterobacterales Recovered From Czech Hospitals, Front Microbiol. 12 (2021) 641415. doi: 10.3389/fmicb.2021.641415.

[45] Y. Sugawara, Y. Akeda, H. Hagiya, N. Sakamoto, D. Takeuchi, R.K. Shanmugakani, et al., Spreading Patterns of NDM-Producing Enterobacteriaceae in Clinical and Environmental Settings in Yangon, Myanmar, Antimicrob Agents Chemother. 63 (2019). doi: 10.1128/aac.01924-18.

[46] P. Lund, A. Tramonti, D. De Biase, Coping with low pH: molecular strategies in neutralophilic bacteria, FEMS Microbiol Rev. 38 (2014) 1091–125. doi: 10.1111/1574-6976.12076.

[47] T.J. Johnson, Y.M. Wannemuehler, L.K. Nolan, Evolution of the iss gene in Escherichia coli, Appl Environ Microbiol. 74 (2008) 2360–9. doi: 10.1128/aem.02634-07.

[48] I.F.N. Lima, N. Boisen, J.D.Q. Silva, A. Havt, E.B. de Carvalho, A.M. Soares, et al., Prevalence of enteroaggregative Escherichia coli and its virulence-related genes in a case-control study among children from north-eastern Brazil, J Med Microbiol. 62 (2013) 683–93. doi: 10.1099/jmm.0.054262-0.

[49] V. Ageorges, R. Monteiro, S. Leroy, C.M. Burgess, M. Pizza, F. Chaucheyras-Durand, et al., Molecular determinants of surface colonisation in diarrhoeagenic Escherichia coli (DEC): from bacterial adhesion to biofilm formation, FEMS Microbiol Rev. 44 (2020) 314–50. doi: 10.1093/femsre/fuaa008.

[50] P. Pramoonjago, M. Kaneko, T. Kinoshita, E. Ohtsubo, J. Takeda, K.S. Hong, et al., Role of TraT protein, an anticomplementary protein produced in Escherichia coli by R100 factor, in serum resistance, J Immunol. 148 (1992) 827–36.

[51] A. Garénaux, M. Caza, C.M. Dozois, The Ins and Outs of siderophore mediated iron uptake by extra-intestinal pathogenic Escherichia coli, Vet Microbiol. 153 (2011) 89–98. doi: 10.1016/j.vetmic.2011.05.023.

[52] N. Boisen, M.T. Østerlund, K.G. Joensen, A.E. Santiago, I. Mandomando, A. Cravioto, et al., Redefining enteroaggregative Escherichia coli (EAEC): Genomic characterization of epidemiological EAEC strains, PLoS Negl Trop Dis. 14 (2020) e0008613. doi: 10.1371/journal.pntd.0008613.

[53] L. Skjøt-Rasmussen, K. Ejrnæs, B. Lundgren, A.M. Hammerum, N. Frimodt-Møller, Virulence factors and phylogenetic grouping of Escherichia coli isolates from patients with bacteraemia of urinary tract origin relate to sex and hospital-vs. community-acquired origin, Int J Med Microbiol. 302 (2012) 129–34. doi: 10.1016/j.ijmm.2012.03.002.

[54] K.E. Rodriguez-Siek, C.W. Giddings, C. Doetkott, T.J. Johnson, M.K. Fakhr, L.K. Nolan, Comparison of Escherichia coli isolates implicated in human urinary tract infection and avian colibacillosis, Microbiology (Reading). 151 (2005) 2097–110. doi: 10.1099/mic.0.27499-0.

[55] S. Yamamoto, T. Tsukamoto, A. Terai, H. Kurazono, Y. Takeda, O. Yoshida, Genetic evidence supporting the fecal-perineal-urethral hypothesis in cystitis caused by Escherichia coli, J Urol. 157 (1997) 1127–9.

[56] L. Li, Y. Zhang, H. Guo, J. Yang, F. He, Genomic insights into a bla(NDM-5)-carrying Escherichia coli ST167 isolate recovered from faecal sample of a healthy individual in China, J Glob Antimicrob Resist. 36 (2024) 240–3. doi: 10.1016/j.jgar.2023.12.032.

[57] Y. Liu, H. Zhang, X. Zhang, N. Jiang, Z. Zhang, J. Zhang, et al., Characterization of an NDM-19-producing Klebsiella pneumoniae strain harboring 2 resistance plasmids from China, Diagn Microbiol Infect Dis. 93 (2019) 355–61. doi: 10.1016/j.diagmicrobio.2018.11.007.

[58] S. Mancini, P.M. Keller, M. Greiner, V. Bruderer, F. Imkamp, Detection of NDM-19, a novel variant of the New Delhi metallo-β-lactamase with increased carbapenemase activity under zinc-limited conditions, in Switzerland, Diagn Microbiol Infect Dis. 95 (2019) 114851. doi: 10.1016/j.diagmicrobio.2019.06.003.

[59] H.A. Hammad, R. Abdelwahab, D.F. Browning, S.A. Aly, Genome Characterization of Carbapenem-Resistant Hypervirulent Klebsiella pneumoniae Strains, Carrying Hybrid Resistance-Virulence IncHI1B/FIB Plasmids, Isolated from an Egyptian Pediatric ICU, Microorganisms. 13 (2025). doi: 10.3390/microorganisms13051058.

[60] A.R. Bizri, A.A. El-Fattah, H.M. Bazaraa, J.W. Al Ramahi, M. Matar, R.A.N. Ali, et al., Antimicrobial resistance landscape and COVID-19 impact in Egypt, Iraq, Jordan, and Lebanon: A survey-based study and expert opinion, PLoS One. 18 (2023) e0288550. doi: 10.1371/journal.pone.0288550.

[61] A. El-Kholy, H.A. El-Mahallawy, N. Elsharnouby, M. Abdel Aziz, A.M. Helmy, R. Kotb, Landscape of Multidrug-Resistant Gram-Negative Infections in Egypt: Survey and Literature Review, Infect Drug Resist. 14 (2021) 1905–20. doi: 10.2147/idr.S298920.

[62] E.A. Scicluna, M.A. Borg, D. Gür, O. Rasslan, I. Taher, S.B. Redjeb, et al., Self-medication with antibiotics in the ambulatory care setting within the Euro-Mediterranean region; results from the ARMed project, J Infect Public Health. 2 (2009) 189–97. doi: 10.1016/j.jiph.2009.09.004.

[63] H. Hafez, M.S. Rakab, A. Elshehaby, A.I. Gebreel, M. Hany, M. BaniAmer, et al., Pharmacies and use of antibiotics: a cross sectional study in 19 Arab countries, Antimicrob Resist Infect Control. 13 (2024) 104. doi: 10.1186/s13756-024-01458-6.

[64] S.W. Elkhadry, M.A.H. Tahoon, Health literacy and its association with antibiotic use and knowledge of antibiotic among Egyptian population: cross sectional study, BMC Public Health. 24 (2024) 2508. doi: 10.1186/s12889-024-19668-3.

[65] K.L. Dooling, A. Kandeel, L.A. Hicks, W. El-Shoubary, K. Fawzi, Y. Kandeel, et al., Understanding Antibiotic Use in Minya District, Egypt: Physician and Pharmacist Prescribing and the Factors Influencing Their Practices, Antibiotics (Basel). 3 (2014) 233–43. doi: 10.3390/antibiotics3020233.

[66] K. Moussally, G. Abu-Sittah, F.G. Gomez, A.A. Fayad, A. Farra, Antimicrobial resistance in the ongoing Gaza war: a silent threat, Lancet. 402 (2023) 1972–3. doi: 10.1016/s0140-6736(23)02508-4.

[67] R. Kumar, O. Tanous, D. Mills, B. Wispelwey, Y. Asi, W. Hammoudeh, et al., Antimicrobial resistance in a protracted war setting: a review of the literature from Palestine, mSystems. 10 (2025) e0167924. doi: 10.1128/msystems.01679-24.

[68] N.A. El Aila, K.I.A. El Aish, Six-year antimicrobial resistance patterns of Escherichia coli isolates from different hospitals in Gaza, Palestine, BMC Microbiol. 25 (2025) 559. doi: 10.1186/s12866-025-04335-3.

