## Supplementary Material for "Antimicrobial resistance and comparative genome analysis of high-risk *Escherichia coli* clones isolated from Egyptian children with diarrhoea"

**Edited by DFB on 1/10/2025**

### Supplementary Material

Radwa Abdelwahab [1, 2], Munirah M. Alhammadi [3], Muhammad Yasir [4],
Ehsan A. Hassan [2], Entsar H. Ahmed [2], Nagla H. Abu-Faddan [2],
Enas A. Daef [2], Stephen J. W. Busby [1] and Douglas F. Browning [5]*

[1] Institute of Microbiology and Infection, School of Biosciences, University of Birmingham, Birmingham, B15 2TT, UK.
[2] Faculty of Medicine, Assiut University, Egypt.
[3] Department of Biology, College of Science, Princess Nourah bint Abdulrahman University, P.O. Box 84428, Riyadh 11671, Saudi Arabia.
[4] Quadram Institute Bioscience, Norwich Research Park, Norwich, NR4 7UQ, UK.
[5] College of Health and Life Sciences, Aston University, Aston Triangle, Birmingham, B4 7ET, UK.

**Table S1.** Antimicrobial susceptibility profile of the MDR *E. coli* strains isolated from children with diarrhoea at Assiut Children's Hospital, Egypt.

| | Antibiotic [a,b] | | | | | | | | | | | |
|---|---|---|---|---|---|---|---|---|---|---|---|---|
| **Strain** | **Ipm** | **Mem** | **Cfc** | **Cro** | **Amc** | **Amp** | **Cip** | **Nor** | **Tob** | **Amk** | **Otc** | **Sxt** |
| **E4** | + | + | + | + | + | + | + | +/- | +/- | +/- | + | + |
| **E15** | + | + | + | + | + | + | + | + | + | - | + | + |
| **E23** | + | + | + | + | + | + | + | + | + | +/- | + | - |
| **E27** | + | + | + | + | + | + | + | + | + | +/- | + | + |
| **E28** | + | + | + | + | + | + | + | + | + | + | + | + |
| **E29** | +/- | +/- | + | + | + | + | + | + | + | + | + | + |
| **E30** | +/- | +/- | + | + | + | + | + | + | + | + | + | + |
| **E34** | + | +/- | + | + | + | + | + | + | + | + | + | + |
| **E35** | + | + | + | + | + | + | + | + | + | + | + | + |
| **E43** | +/- | +/- | + | + | +/- | + | + | + | + | + | + | + |

[a] Antibiotics used were as follows: Ipm, imipenem; Mem, meropenem; Cfc, cefaclor; Cro, ceftriaxone; Amc, amoxicillin; Amp, ampicillin; Cip, ciprofloxacin; Nor, norfloxacin; Tob, tobramycin; Amk, amikacin; Otc, oxytetracycline; Sxt, trimethoprim/ sulfamethoxazole.
[b] Antibiotic resistant (+), susceptible (-) and intermediate (+/-), *i.e.* above the point of susceptibility but below the resistant breakpoint [1].

**Table S2.** Analysis of chromosomal point mutations associated with nalidixic acid and ciprofloxacin resistance carried by the *E. coli* strains used in this study.

| **Strain** | **Chromosomal point mutations associated with AMR [a]** | **ST [b]** |
|---|---|---|
| **E4** | N/D | ST46 |
| **E15** | *gyrA* S83L D87N , *parC* S80I, *parE* S458A | ST167 |
| **E23** | *gyrA* S83L D87N, *parC* S80I, *parE* S458A | ST167 |
| **E27** | *gyrA* S83L D87N, *parC* S80I, *parE* S458A | ST410 |
| **E28** | *gyrA* S83L D87N, *parC* S80I, *parE* S458A | ST617 |
| **E29** | *gyrA* S83L, D87N, *parC* S80I E84G | ST361 |
| **E30** | *gyrA* S83L D87N, *parC* S80I, *parE* S458A | ST410 |
| **E34** | *gyrA* S83L D87N, *parC* S80I, *parE* S458A | ST410 |
| **E35** | *gyrA* S83L D87N, *parC* S80I, *parE* S458A | ST167 |
| **E43** | *gyrA* S83L D87N, *parC* S80I, *parE* S458A | ST167 |

Software at the Center for Genomic Epidemiology (CGE) (http://www.genomicepidemiology.org/ (accessed on 10th July 2025)) was used to identify: [a] the chromosomal point mutations associated with nalidixic acid and ciprofloxacin resistance [2] and [b] the sequence type (ST) [3] of each strain. N/D none detected.

**Table S3.** Analysis of the potential virulence genes carried by the *E. coli* strains used in this study.

| **Strain** | **ST** [a] | **Virulence determinants** [b] | **Pathogen Finder Score** [c] |
|---|---|---|---|
| **E4** | ST46 | *gad, iss, terC* | 0.929 |
| **E15** | ST167 | *capU, gad, hra, iss, terC, traT,* ***fyuA***, ***irp2*** | 0.925 |
| **E23** | ST167 | *capU, gad, hra, iss, terC, traT* | 0.928 |
| **E27** | ST410 | *gad, lpfA, terC* | 0.937 |
| **E28** | ST617 | *gad, iss, sitA, terC, traT,* ***fyuA***, ***irp2***, ***iucC***, ***iutA*** | 0.928 |
| **E29** | ST361 | *astA, capU, gad, sitA, terC, traT* | 0.93 |
| **E30** | ST410 | *gad, lpfA, terC* | 0.939 |
| **E34** | ST410 | *gad, lpfA, terC* | 0.937 |
| **E35** | ST167 | *capU, gad, hra, iss, terC, traT* | 0.92 |
| **E43** | ST167 | *capU, gad, hra, iss, terC, traT* | 0.927 |

Software at the CGE was used to identify: [a] the sequence type (ST) [3] of each strain, [b] the potential virulence genes they carry [4] and [c] if they were likely a human pathogen using PathogenFinder 1.1 [5]. For PathogenFinder scores range from 0 to 1 with values closer to 1 indicating that the input organism was predicted as a human pathogen.

**Supplementary Figure legends**

**Figure S1.** Comparison of *E. coli* plasmid pLAU-NDM19 with the draft genome of *E. coli* E23. **A)** The panel shows the comparison of pLAU-NDM19 (CP074195.1) [6] with the draft genome of E23 and E23 contig 27 (46,073 bp) using ProkSee [7]. The outer two rings display the genes of pLAU-NDM19 (CDS) on both strands. The green and brown rings illustrate the BLAST results when the E23 draft genome and contig 27, respectively, are compared to pLAU-NDM19. **B)** The panel shows the alignment pLAU-NDM19 (CP074195.1) [6] with E23 contig 27 (46,073 bp) using ACT [8]. Alignment is shown by blue banding, which indicates that the sequences are inverted with respect to each other. The location of the $\textit{bla}_{\text{NDM-19}}$ in both panels is indicated. **C)** The panel shows the alignment of the amino acid sequences of the NDM-1 (AHM26723), NDM-5 (JN104597), NDM-7 (AKN35289) and NDM-19 (WP_094009810.1) carbapenemases with the NDM-19 carbapenemase carried by *E. coli* strain E23. Differences from NDM-1 are highlighted red.

**Figure S2.** Comparison of *E. coli* plasmid pEcMAD2 with the draft genome of *E. coli* E27. **A)** The panel shows the comparison of pEcMAD2 (LR595693.1) [9] with the draft genome of E27, E27 contig 22 (48,979 bp), plasmid pE2-OXA-181 (CP048918.1) [10] and plasmid pEc1079_3 (CP081309.1) [11],using ProkSee [7]. The outer rings display the genes of pEcMAD2 (CDS) on both strands. The green, light green, brown and blue rings illustrate the BLAST results when the E27 draft genome, E27 contig 22, pE2-OXA-181 and pEc1079_3 are compared to pEcMAD2. **B)** The panel shows the alignment of pEcMAD2 (LR595693.1) [9] with E27 contig 22 (48,979 bp) using ACT [8]. Alignment is shown by red banding. The location of the $\textit{bla}_{\text{OXA-181}}$ and the IncX3 and ColKP3 replicons panels is indicated in both.

**Figure S3.** Comparison of *E. coli* EcMAD1 and E2 chromosomes with the draft genomes of *E. coli* E27, E30 and E34. **A)** The panel shows the comparison of the EcMAD1 chromosome (LR595691.1: 4,747,851 bp) [9] with the draft genomes of E27, E30 and E34, and the chromosomes of *E. coli* strains E2 (CP048915.1) [10] and Ec1079 (CP081306.1) [11], using ProkSee [7]. The outer rings display the genes of EcMAD1 (CDS) on both strands. The green, brown, purple, orange and blue rings illustrate the BLAST results when the E27, E30 and E34 draft genomes and the chromosomes of *E. coli* strains E2 and Ec1079 are compared to EcMAD1. **B)** The panel shows the comparison of the E2 chromosome (CP048915.1: 4,741,120 bp) [10] with the draft genomes of E27, E30, E34, and the chromosomes of *E. coli* strains EcMAD1 (LR595691.1 bp) [9] and Ec1079 (CP081306.1) [11], using ProkSee [7]. The outer rings display the genes of E2 (CDS) on both strands. The green, brown, purple, orange and blue rings illustrate the BLAST results when the E27, E30 and E34 draft genomes, and the chromosomes of *E. coli* strains EcMAD1 and Ec1079 are compared to E2.

**Figure S4.** Comparison of plasmid pM309-NDM5 with the draft genome of stain E35. **A)** The panel shows the comparison of pM309-NDM5 (AP018833.1: 136,947 bp) [12] with the draft genome of E35 and E35 contigs 38 (38,086 bp), 45 (IncFIA replicon: 15,794 bp) and 49 (IncFII replicon: 11,728 bp), using ProkSee [7]. The genes (CDS) of pM309-NDM5 are displayed in the outer rings, with the location of the various AMR genes (including $bla_{\text{NDM-5}}$) and plasmid replicons (IncFIA and IncFII) indicated. The green, brown, light brown and light green rings depict the BLAST results when the E35 draft genome and contigs 38, 45 and 49 are compared with pM309-NDM5. **B)** The panel shows the alignment of pM309-NDM5 (AP018833.1) [12] with E35 contig 38 (38,086 bp) using ACT [8]. Alignment is shown by blue banding, which indicates that the sequences are inverted with respect to each other. The location of the $bla_{\text{NDM-5}}$ is indicated.

**Figure S5.** Analysis of contigs 22 and 36 from *E. coli* strain E28. Genomic organisation of E28 **A)** contig 36 (43,206 bp) and **B)** contig 22 (75,303 bp) using ProkSee [7]. **C)** Comparison of plasmid pEC22-OXA-1 from *E. coli* strain Ec20 with the draft genome of *E. coli* E28. The panel shows the comparison of pEC22-OXA-1 (CP084902.1: 169208 bp) with the draft genome of E28, E28 contigs 22 (75,303 bp) and 36 (43,206 bp), and plasmid pEC22-CTX-M-15 (CP157417.1: 169,208 bp), using ProkSee [7]. The outer two rings display the genes of pEC22-OXA-1 (CDS) on both strands. The green, brown, light brown and purple rings illustrate the BLAST results when the E28 draft genome, contigs 22 and 36, and pEC22-CTX-M-15, respectively, are compared to pEC22-OXA-1. **D)** The panel shows the alignment of pEC22-OXA-1 (CP084902.1: 169,208 bp) with E28 contigs 22 (75,303 bp) and 36 (43,206 bp) using ACT [8]. Alignment is shown by red and blue banding.

**Figure S6.** Comparison of *E. coli* plasmid pE2-2 with the draft genomes of *E. coli* E27 and E34. **A)** The panel shows the comparison of plasmid pE2-2 (CP048917.1: 92,027 bp) [10] with the with the draft genome and contig 28 (92,136 bp) of E27, and the draft genome of E34 and contig 16 (96,506 bp), using ProkSee [7]. The outer rings display the CDS of pE2-2 on both strands. The green, light green, brown and light brown rings illustrate the BLAST results when the E27 draft genome, E27 contig 18, E34 draft genome and E34 contig 16, respectively, are compared to pE2-2. **B)** The panel shows the alignment of E34 contig 16 (96,506 bp), E27 contig 18 (92,136 bp) with pE2-2 (CP048917.1) [10], using ACT [8]. Alignment is shown by red and blue banding. Blue banding indicates that the sequences have been inverted with respect to each other. The location of the p0111 replicon is indicated in both panels.

**Figure S7.** Analysis of *E. coli* plasmid pEcMAD1. The panel shows the comparison of plasmid pEcMAD1 (LR595692.1: 98,473 bp) [9] with the with the draft genomes of E27, E30 and E34 and plasmids pE2-

NDM-CTX-M (CP048916.1) [10] and pEc1079_1 (CP081307.1) [11], using ProkSee [7]. The outer rings display the CDS of pEcMAD1 on both strands. The green, brown, purple, orange and blue rings illustrate the BLAST results when pEcMAD1 is compared to the draft genomes of E27, E30 and E34 and pE2-NDM-CTX-M and pEc1079_1.

**Supplementary References**

[1] Clinical_and_Laboratory_Standards_Institute_(CLSI), Performance Standards for Antimicrobial Susceptibility Testing; Twenty-Fourth Informational Supplement, CLSI Document M100-S24, Wayne. 34(1) (2014).
[2] E. Zankari, H. Hasman, S. Cosentino, M. Vestergaard, S. Rasmussen, O. Lund, et al., Identification of acquired antimicrobial resistance genes, J Antimicrob Chemother. 67 (2012) 2640-4. doi: 10.1093/jac/dks261.
[3] M.V. Larsen, S. Cosentino, S. Rasmussen, C. Friis, H. Hasman, R.L. Marvig, et al., Multilocus sequence typing of total-genome-sequenced bacteria, J Clin Microbiol. 50 (2012) 1355-61. doi: 10.1128/jcm.06094-11.
[4] K.G. Joensen, F. Scheutz, O. Lund, H. Hasman, R.S. Kaas, E.M. Nielsen, et al., Real-time whole-genome sequencing for routine typing, surveillance, and outbreak detection of verotoxigenic *Escherichia coli*, J Clin Microbiol. 52 (2014) 1501-10. doi: 10.1128/jcm.03617-13.
[5] S. Cosentino, M. Voldby Larsen, F. Møller Aarestrup, O. Lund, PathogenFinder--distinguishing friend from foe using bacterial whole genome sequence data, PLoS One. 8 (2013) e77302. doi: 10.1371/journal.pone.0077302.
[6] J. Moussa, E. Nassour, T. Jisr, M. El Chaar, S. Tokajian, Characterization of bla(NDM-19)-producing IncX3 plasmid isolated from carbapenem-resistant *Escherichia coli* and *Klebsiella pneumoniae*, Heliyon. 10 (2024) e29642. doi: 10.1016/j.heliyon.2024.e29642.
[7] J.R. Grant, E. Enns, E. Marinier, A. Mandal, E.K. Herman, C.Y. Chen, et al., Proksee: in-depth characterization and visualization of bacterial genomes, Nucleic Acids Res. 51 (2023) W484-w92. doi: 10.1093/nar/gkad326.
[8] T.J. Carver, K.M. Rutherford, M. Berriman, M.A. Rajandream, B.G. Barrell, J. Parkhill, ACT: the Artemis Comparison Tool, Bioinformatics. 21 (2005) 3422-3. doi: 10.1093/bioinformatics/bti553.
[9] R. Patiño-Navarrete, I. Rosinski-Chupin, N. Cabanel, L. Gauthier, J. Takissian, J.Y. Madec, et al., Stepwise evolution and convergent recombination underlie the global dissemination of carbapenemase-producing *Escherichia coli*, Genome Med. 12 (2020) 10. doi: 10.1186/s13073-019-0699-6.
[10] D. Gamal, M. Fernández-Martínez, I. El-Defrawy, A.A. Ocampo-Sosa, L. Martínez-Martínez, First identification of NDM-5 associated with OXA-181 in Escherichia coli from Egypt, Emerg Microbes Infect. 5 (2016) e30. doi: 10.1038/emi.2016.24.
[11] S. Mahazu, I. Prah, A. Ayibieke, W. Sato, T. Hayashi, T. Suzuki, et al., Possible Dissemination of *Escherichia coli* Sequence Type 410 Closely Related to B4/H24RxC in Ghana, Front Microbiol. 12 (2021) 770130. doi: 10.3389/fmicb.2021.770130.
[12] Y. Sugawara, Y. Akeda, H. Hagiya, N. Sakamoto, D. Takeuchi, R.K. Shanmugakani, et al., Spreading Patterns of NDM-Producing Enterobacteriaceae in Clinical and Environmental Settings in Yangon, Myanmar, Antimicrob Agents Chemother. 63 (2019). doi: 10.1128/aac.01924-18.

**Supplementary Figure S1.**

**(A)**

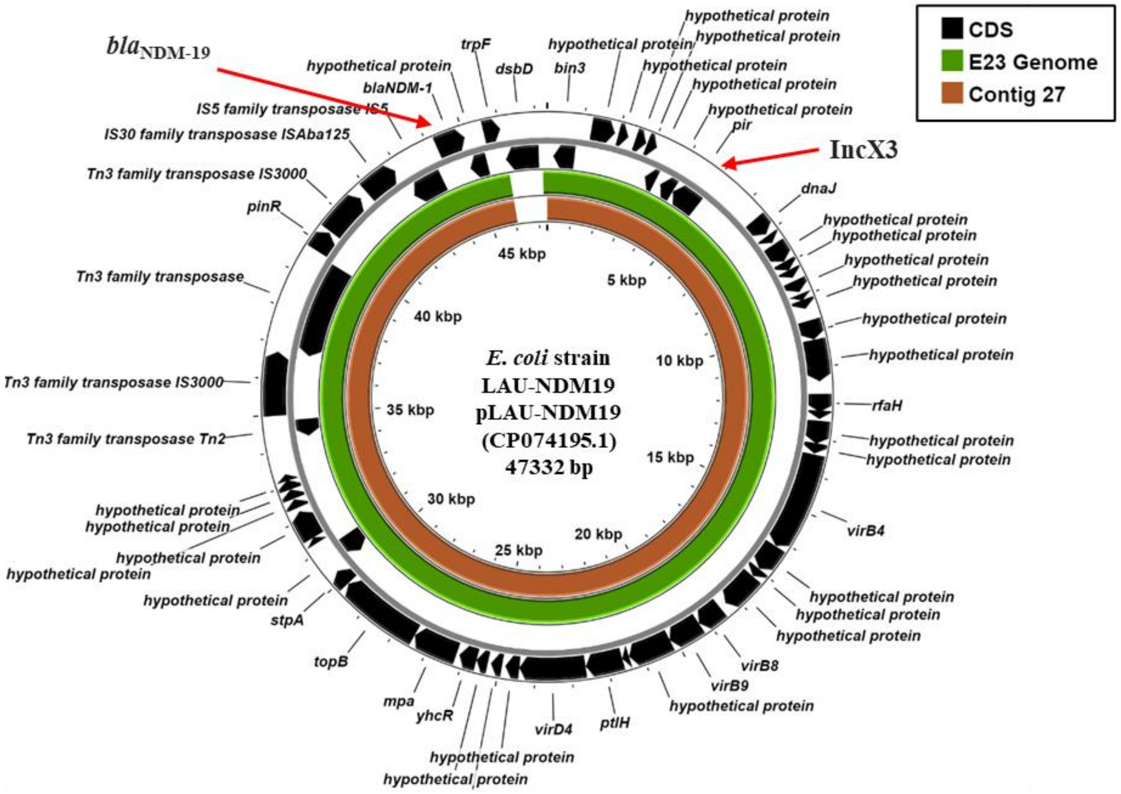


**(B)**

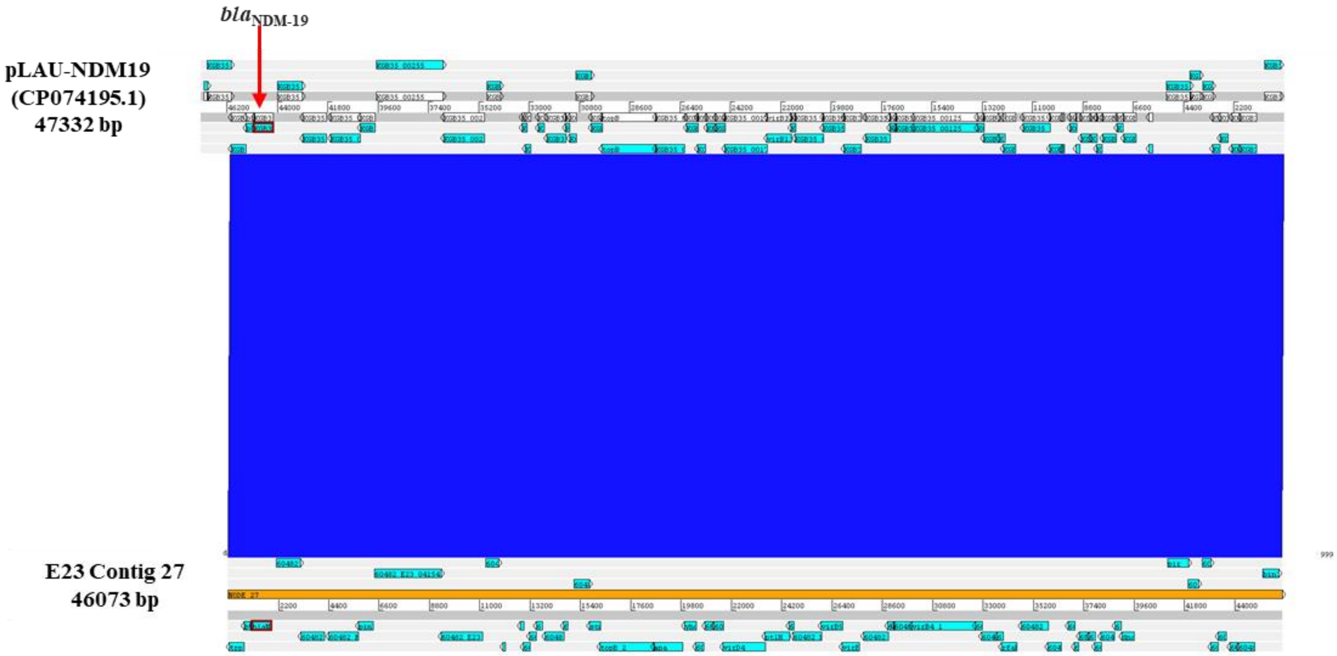

**Supplementary Figure S1 (continued).**

**(C)**

```
NDM-19_E23      MELPNIMHPVAKLSTALAAALMLSGCMPGEIRPTIGQQMETGDQRFGDLVFRQLAPNVWQ
NDM-19          MELPNIMHPVAKLSTALAAALMLSGCMPGEIRPTIGQQMETGDQRFGDLVFRQLAPNVWQ
NDM-7           MELPNIMHPVAKLSTALAAALMLSGCMPGEIRPTIGQQMETGDQRFGDLVFRQLAPNVWQ
NDM-1           MELPNIMHPVAKLSTALAAALMLSGCMPGEIRPTIGQQMETGDQRFGDLVFRQLAPNVWQ
NDM-5           MELPNIMHPVAKLSTALAAALMLSGCMPGEIRPTIGQQMETGDQRFGDLVFRQLAPNVWQ
                ************************************************************

NDM-19_E23      HTSYLDMPGFGAVASNGLIVRDGGRVLVVDTAWTDDQTAQILNWIKQEINLPVALAVVTH
NDM-19          HTSYLDMPGFGAVASNGLIVRDGGRVLVVDTAWTDDQTAQILNWIKQEINLPVALAVVTH
NDM-7           HTSYLDMPGFGAVASNGLIVRDGGRVLVVDTAWTDDQTAQILNWIKQEINLPVALAVVTH
NDM-1           HTSYLDMPGFGAVASNGLIVRDGGRVLVVDTAWTDDQTAQILNWIKQEINLPVALAVVTH
NDM-5           HTSYLDMPGFGAVASNGLIVRDGGRVLLVDTAWTDDQTAQILNWIKQEINLPVALAVVTH
                ***************************:********************************

NDM-19_E23      AHQDKMGGMNALHAAGIATYANALSNQLAPQEGLVAAQHSLTFAANGWVEPATAPNFGPL
NDM-19          AHQDKMGGMNALHAAGIATYANALSNQLAPQEGLVAAQHSLTFAANGWVEPATAPNFGPL
NDM-7           AHQDKMGGMNALHAAGIATYANALSNQLAPQEGLVAAQHSLTFAANGWVEPATAPNFGPL
NDM-1           AHQDKMGGMDALHAAGIATYANALSNQLAPQEGMVAAQHSLTFAANGWVEPATAPNFGPL
NDM-5           AHQDKMGGMDALHAAGIATYANALSNQLAPQEGLVAAQHSLTFAANGWVEPATAPNFGPL
                *********:***********************:**************************

NDM-19_E23      KVFYPGPGHTSDNITVGIDGTDIAFGGCLIKDSKAKSLGNLGDADTEHYAASVRAFGAAF
NDM-19          KVFYPGPGHTSDNITVGIDGTDIAFGGCLIKDSKAKSLGNLGDADTEHYAASVRAFGAAF
NDM-7           KVFYPGPGHTSDNITVGIDGTDIAFGGCLIKDSKAKSLGNLGDADTEHYAASARAFGAAF
NDM-1           KVFYPGPGHTSDNITVGIDGTDIAFGGCLIKDSKAKSLGNLGDADTEHYAASARAFGAAF
NDM-5           KVFYPGPGHTSDNITVGIDGTDIAFGGCLIKDSKAKSLGNLGDADTEHYAASARAFGAAF
                ****************************************************.*******

NDM-19_E23      PKASMIVMSHSAPDSRAAITHTARMADKLR
NDM-19          PKASMIVMSHSAPDSRAAITHTARMADKLR
NDM-7           PKASMIVMSHSAPDSRAAITHTARMADKLR
NDM-1           PKASMIVMSHSAPDSRAAITHTARMADKLR
NDM-5           PKASMIVMSHSAPDSRAAITHTARMADKLR
                ******************************
```

**Supplementary Figure S2.**

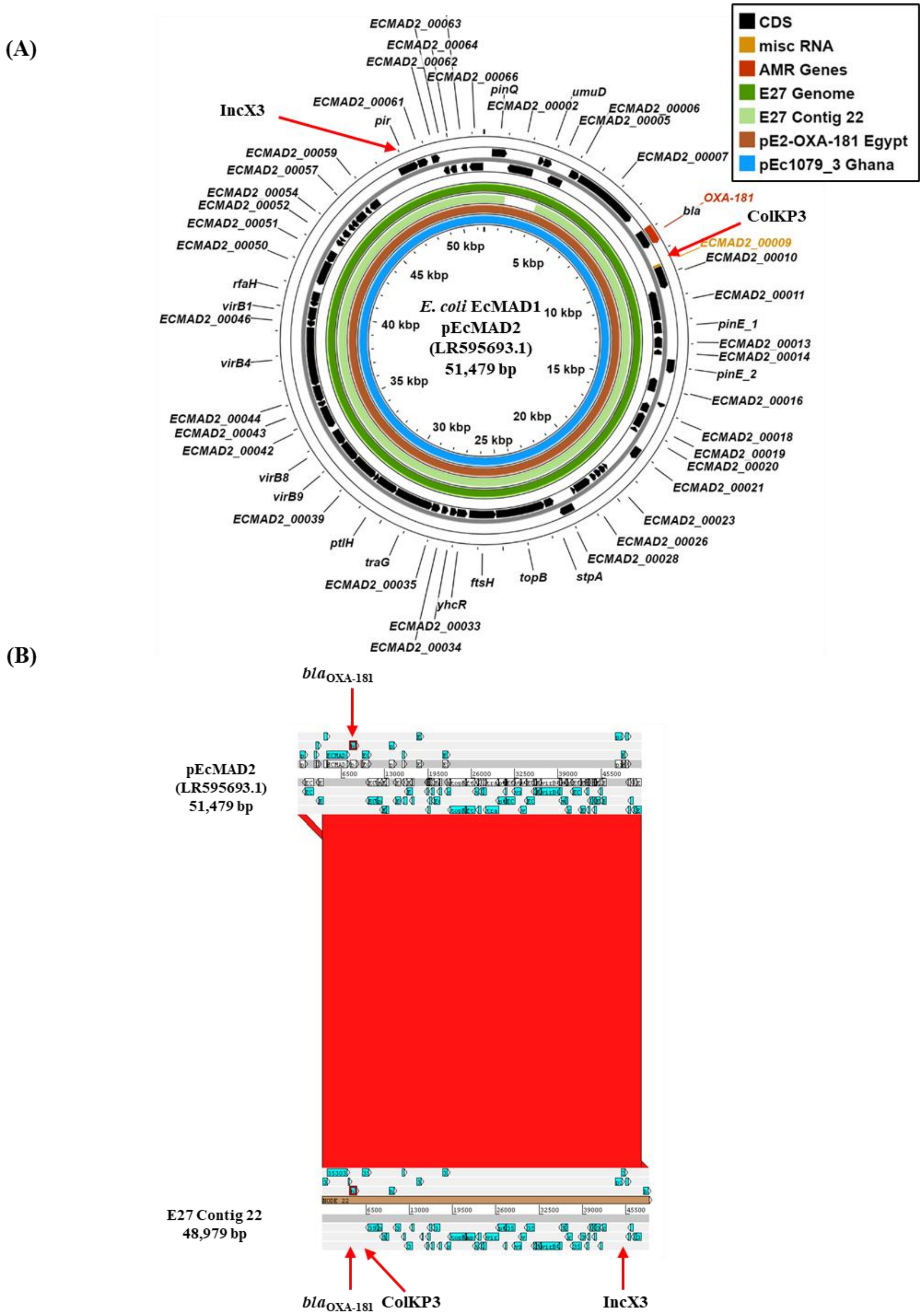

**Supplementary Figure S3.**

**(A)**

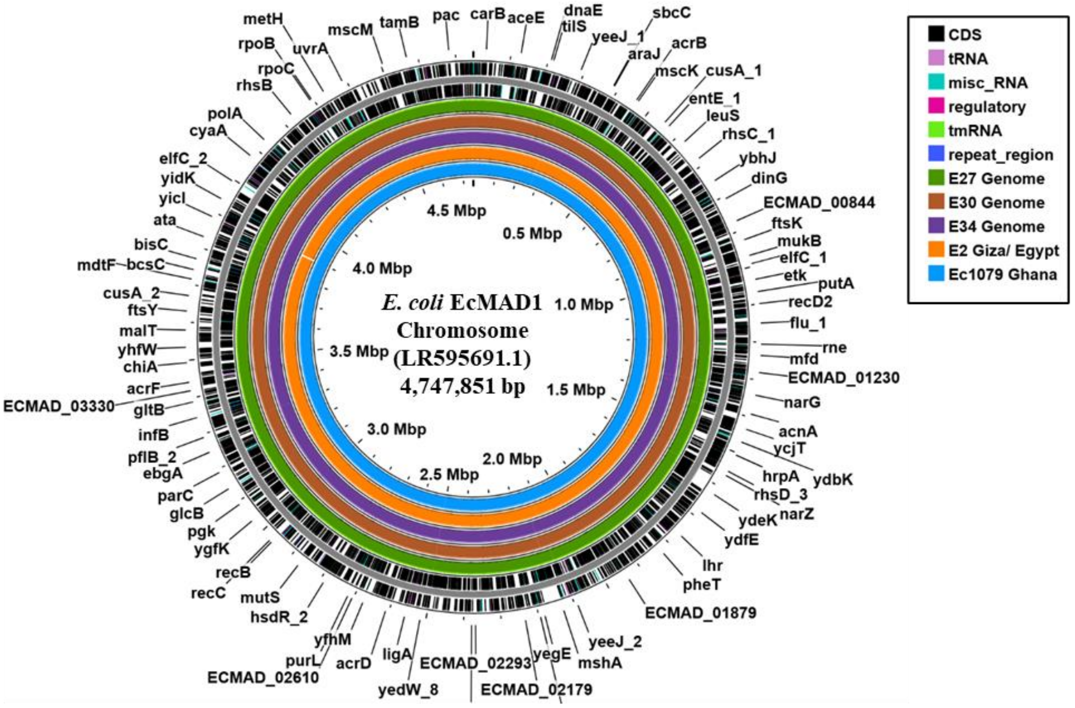


**(B)**

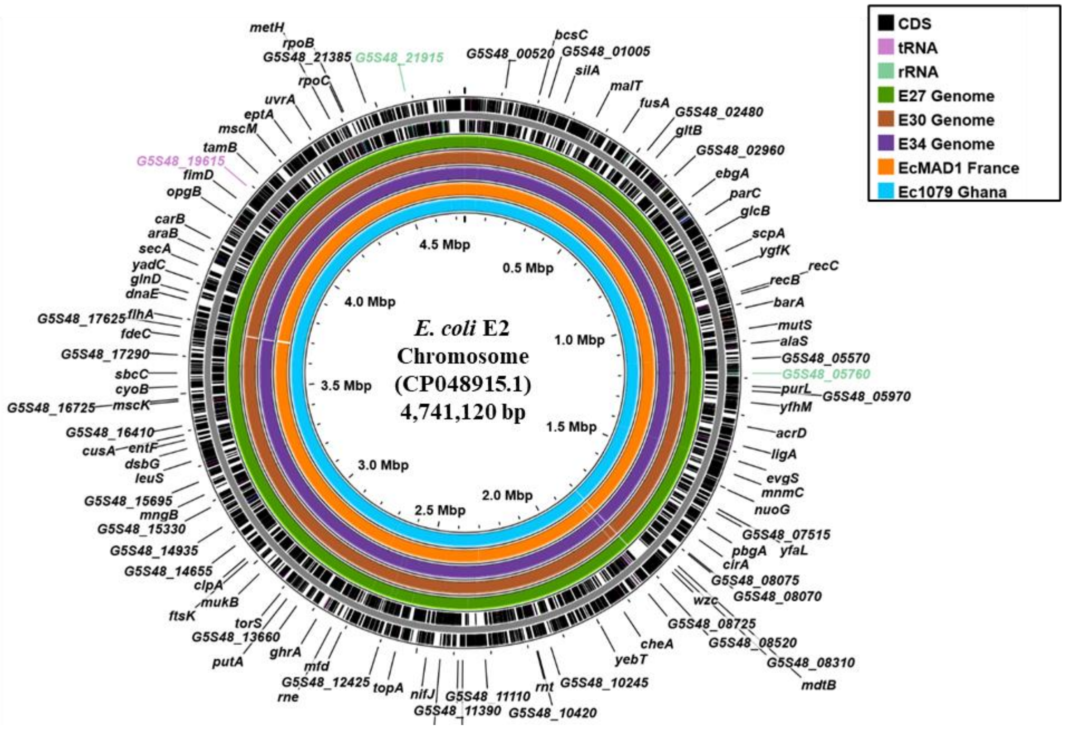

**Supplementary Figure S4.**

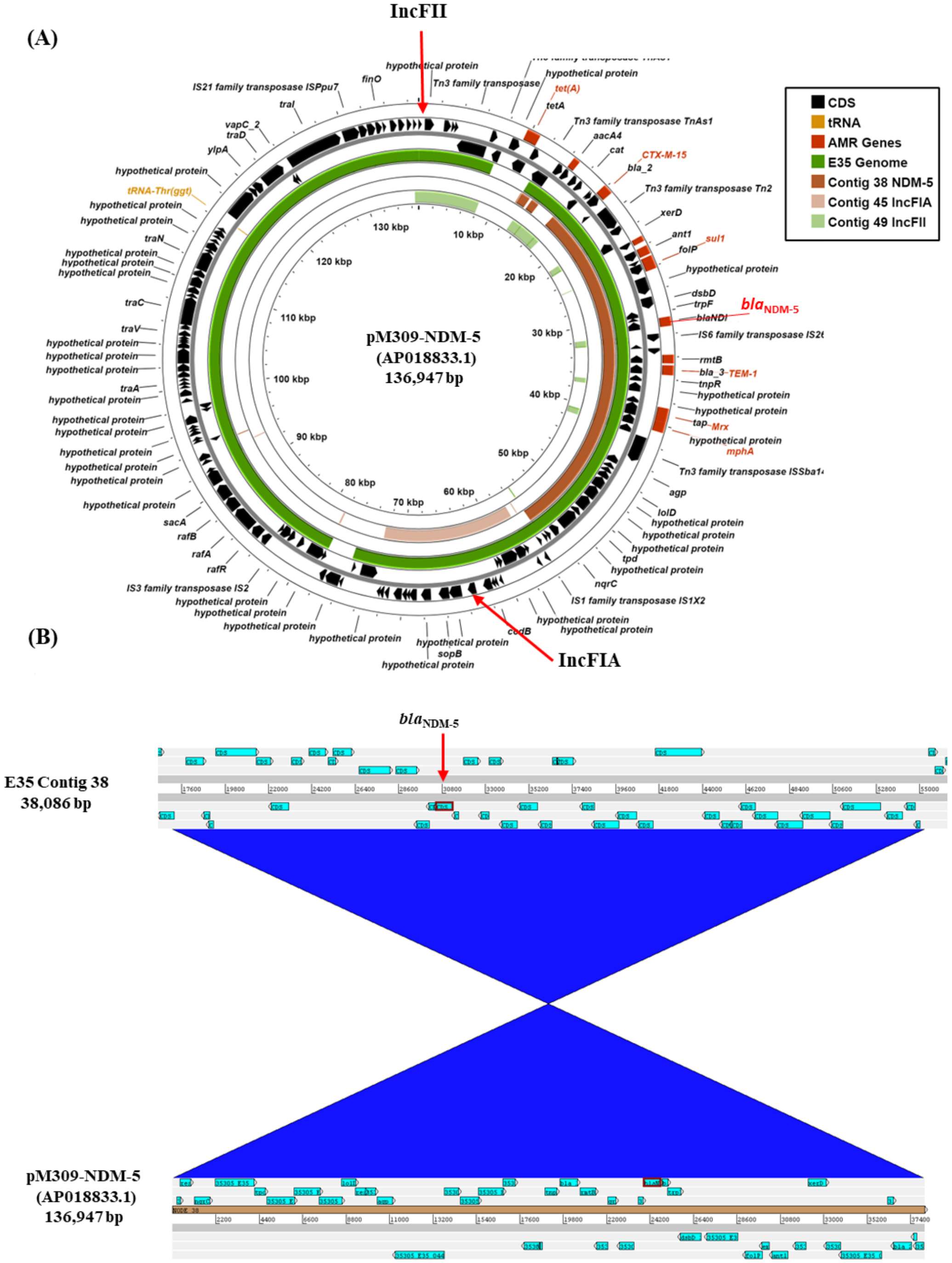

**Supplementary Figure S5.**

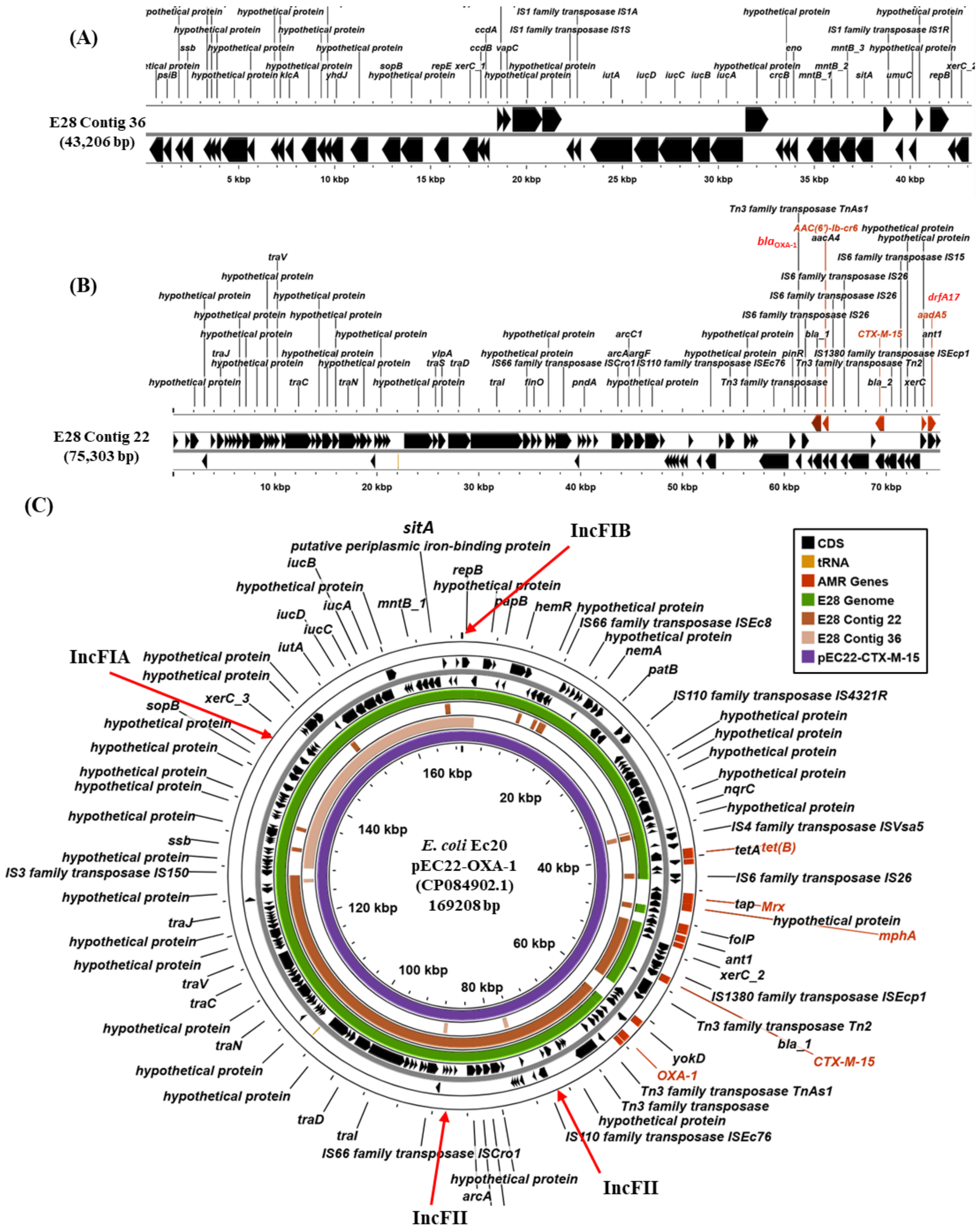

**Supplementary Figure S5. (cont)**

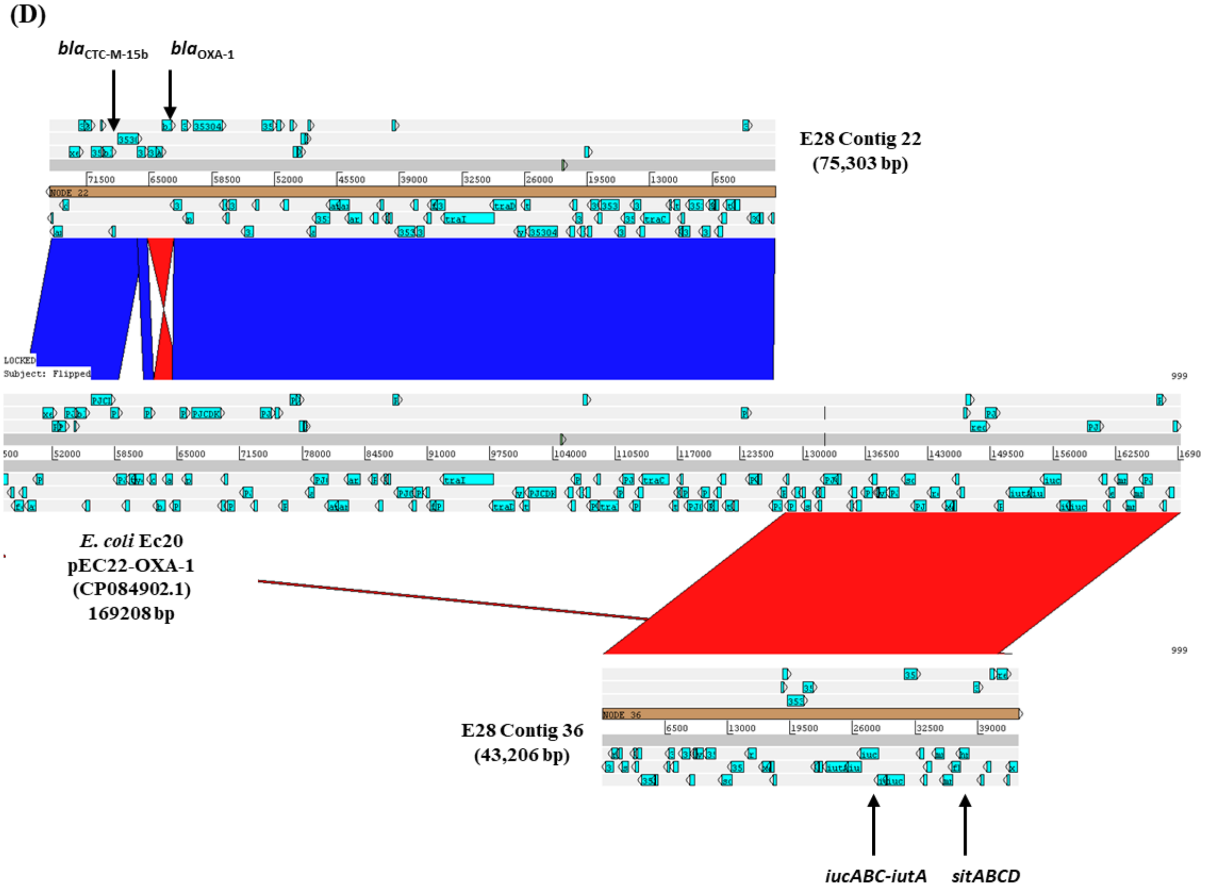

**Supplementary Figure S6.**

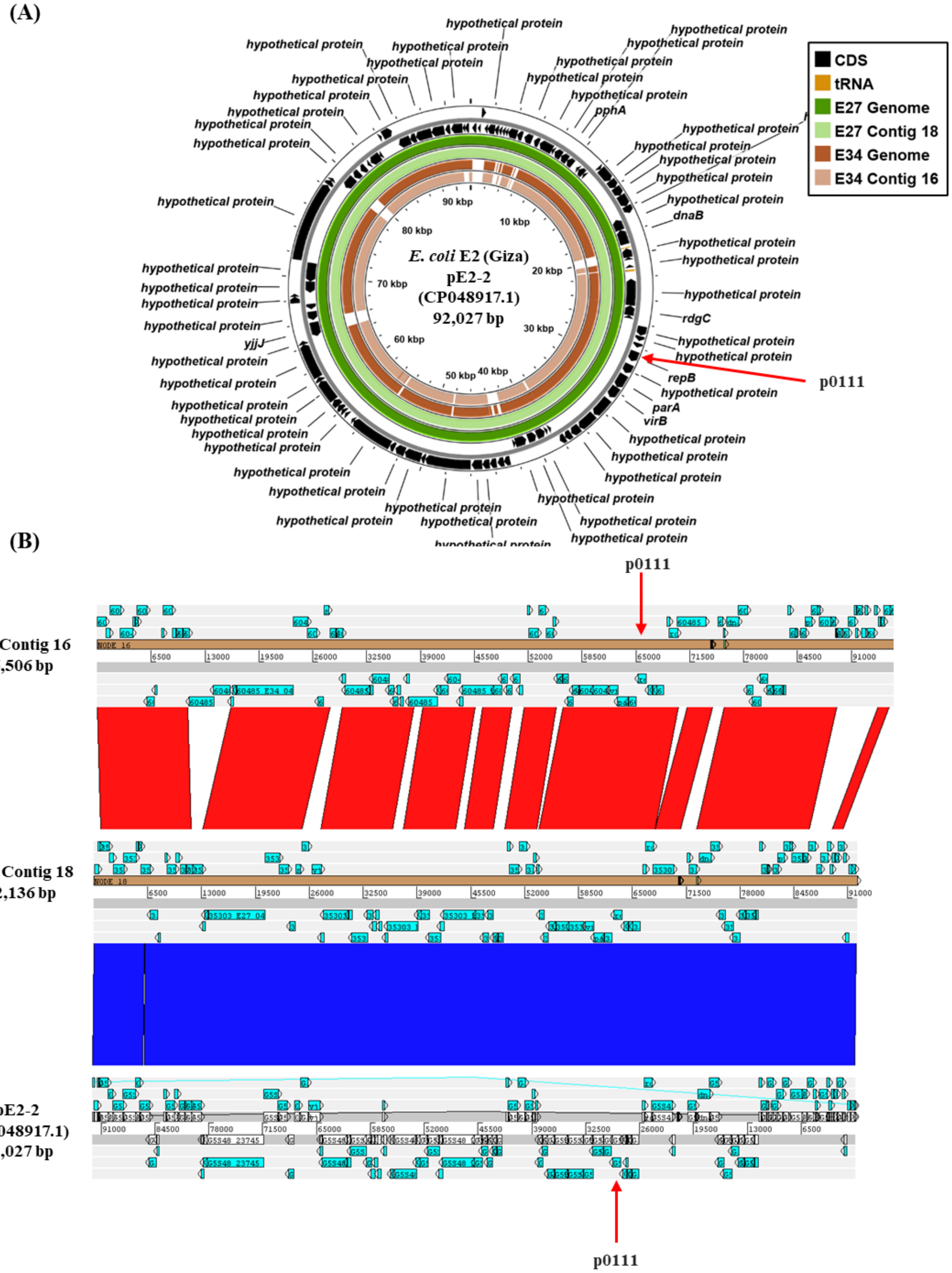

**Supplementary Figure S7.**

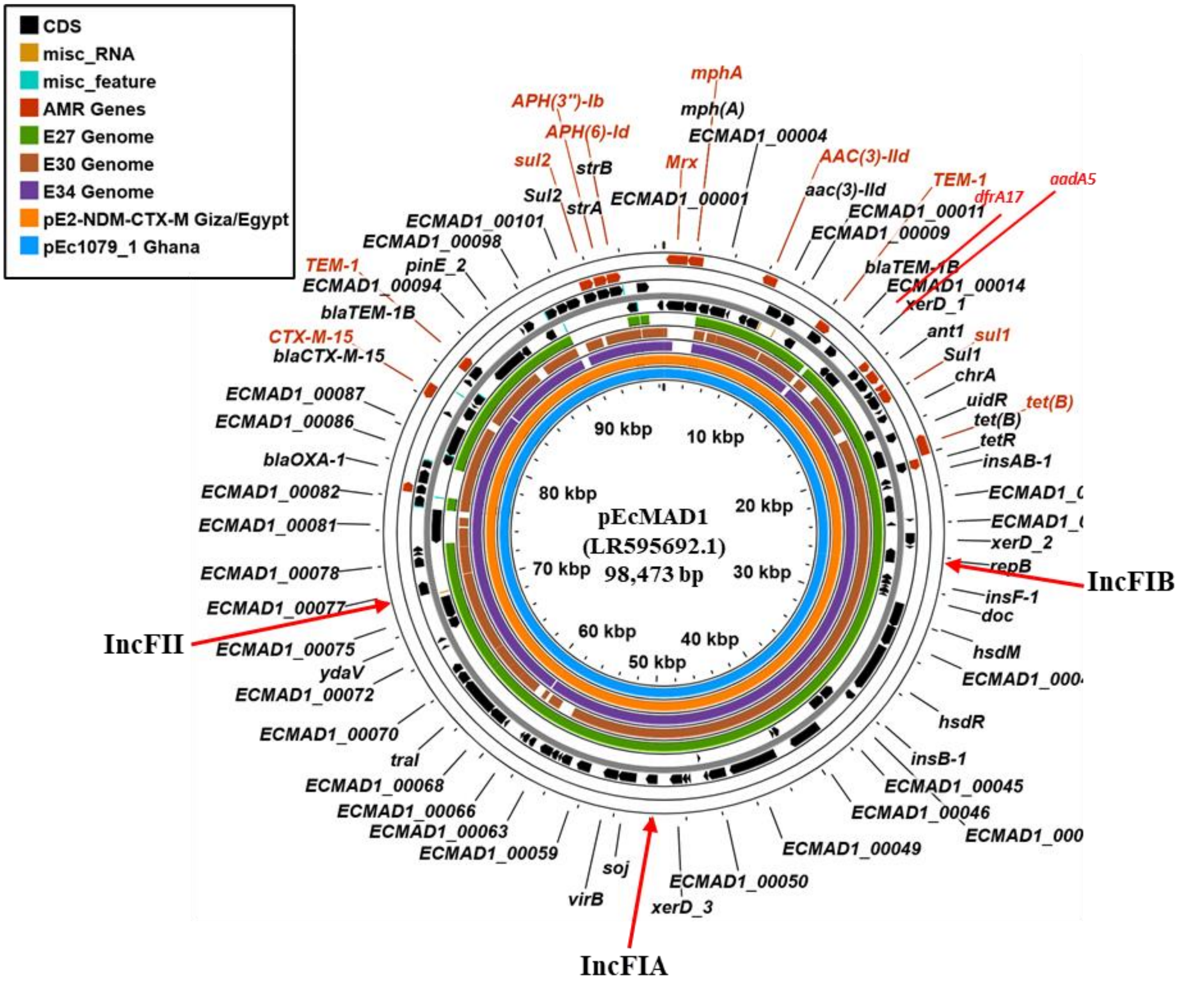
